# Structural basis of bacteriophage T5 infection trigger and *E. coli* cell wall perforation

**DOI:** 10.1101/2022.09.20.507954

**Authors:** Romain Linares, Charles-Adrien Arnaud, Grégory Effantin, Claudine Darnault, Nathan Hugo Epalle, Elisabetta Boeri Erba, Guy Schoehn, Cécile Breyton

## Abstract

The vast majority of bacteriophages (phages) - bacterial viruses - present a tail that allows host recognition, cell wall perforation and safe channelling of the viral DNA from the capsid to the cytoplasm of the infected bacterium. The majority of tailed phages bears a long flexible tail (*Siphoviridae*) at the distal end of which a tip complex, often called baseplate, harbours one or more Receptor Binding Protein·s (RBPs). Interaction between the RBPs and the host surface triggers cell wall perforation and DNA ejection, but little is known on these mechanisms for *Siphoviridae*. Here, we present the structure of siphophage T5 tip at high resolution, determined by electron cryo-microscopy, allowing to trace most of its constituting proteins, including 35 C-terminal residues of the Tape Measure Protein. We also present the structure of T5 tip after interaction with its *E. coli* receptor FhuA reconstituted into nanodisc. It brings out the dramatic conformational changes underwent by T5 tip upon infection, *i*.*e*. bending of the central fibre on the side, opening of the tail tube and its anchoring to the membrane, and formation of a transmembrane channel. These new structures shed light on the mechanisms of host recognition and activation of the viral entry for *Siphoviridae*.

## Introduction

Bacteriophages or phages, viruses that infect bacteria, represent the most abundant biological entity on our planet. They are present in all ecosystems where bacteria develop, and outnumber their hosts by at least an order of magnitude, being instrumental in the development and evolution of microbial populations (1). Moreover, with the increasing number of pathogenic strains resistant to antibiotics, virulent phages are considered as a serious alternative or complement to classical treatments (2). The vast majority of known phages bear a tail whose tip serves to recognise the host, perforate the bacterial cell wall and deliver the viral genome into the host cytoplasm. Tails can be long and contractile in *Myoviridae*, long and flexible in *Siphoviridae* or short in *Podoviridae*. Bacterial tail-like machines also serve as a means to inject various macromolecules in neighbouring prokaryotic and/or eukaryotic cells: all these systems derive from a common ancestor that share high structural similarities (3–6). Whereas the contracting tails and tail-like systems have seen their mechanism of sheath contraction and inner tube propelling described in exquisite details (5, 7–9), very little is known on the mechanisms of host recognition transmission, tail opening and cell wall perforation in Siphophages that represent more than 60% of all phages (10).

Phage T5 (11), a *Siphoviridae* infecting *E. coli*, is a model phage belonging to the T-series introduced by Delbrück and co-workers in the 1940s (12). It presents a 90-nm icosahedral capsid (13) to which is attached a 160-nm tail tube (Fig. 1A), formed by the polymerisation of 40 ring-shaped trimers of the Tail Tube Protein pb6 (TTP_pb6_)(14) around the Tape Measure Protein pb2 (TMP_pb2_)(15). At its distal end, the tail harbours the tip complex, also called baseplate: three dispensable Side Tail Fibres (STF_pb1_) reversibly bind to the sugar moiety of the host lipopolysaccharide (16). They are linked, by the collar to a conical structure formed by the Distal Tail Protein pb9 (DTP_pb9_)(17) and the Baseplate Hub Protein pb3 (BHP_pb3_, also called Tal, Tail Associated Lysin/Lysozyme)(11). A central fibre protein (pb4), at the extremity of which is found the Receptor Binding Protein pb5 (RBP_pb5_)(18, 19), completes the tip complex, with p140 and p132 of unknown location (11)(Fig. 1B). FhuA, an outer-membrane E. coli iron-ferrichrome transporter, is the bacterial receptor recognized by T5 (20). The mere interaction of T5 with its purified receptor FhuA triggers the release of the DNA in vitro (21), making this phage an excellent tool for studying host recognition, DNA ejection (14, 22) and cell wall perforation mechanisms. We thus embarked on solving the structure of T5 tail tip before (Tip) and after (Tip-FhuA) interaction with its receptor to unravel the conformational changes induced and understand the mechanism of tail opening and cell wall perforation in *Siphoviridae*.

**Figure 1:**
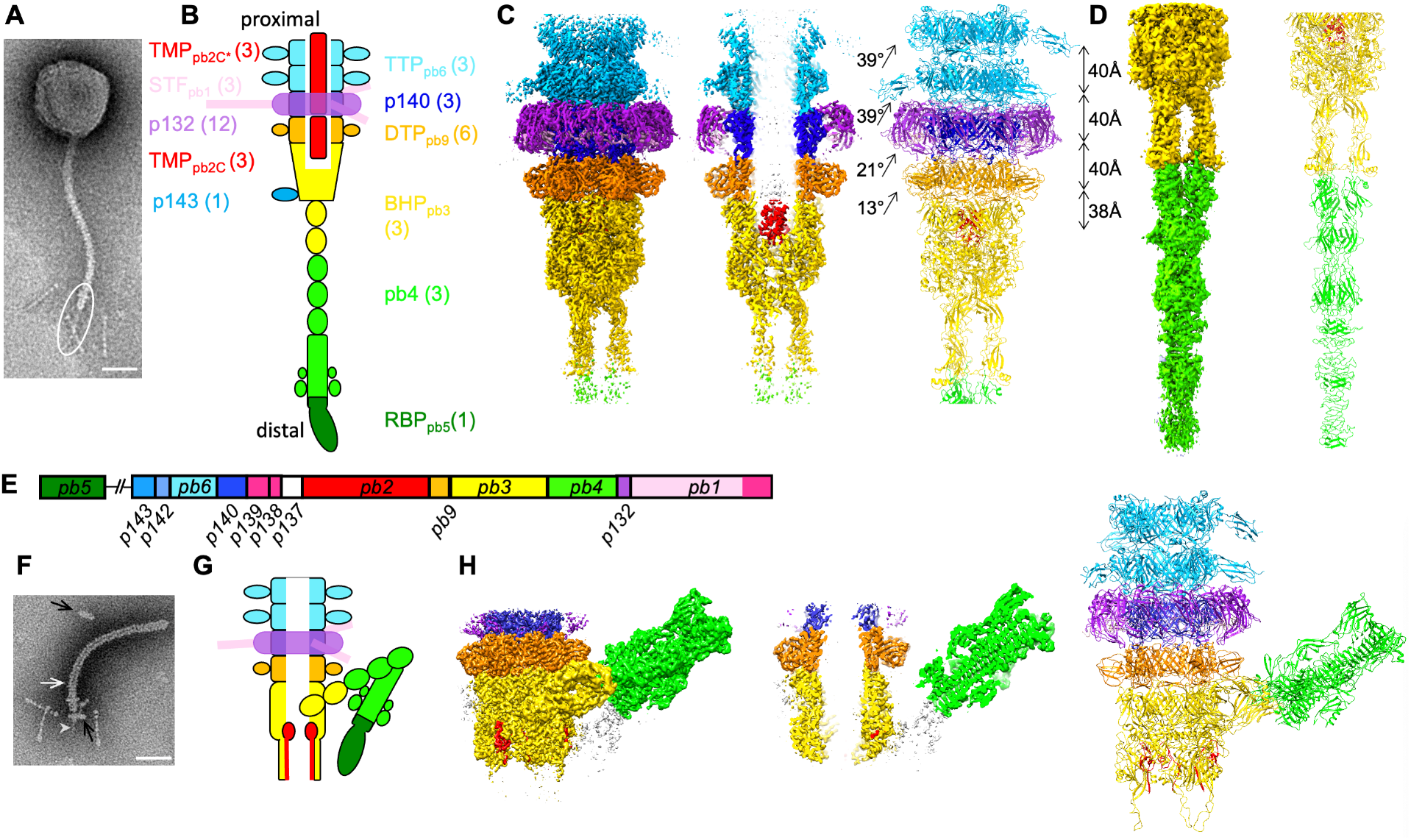
Structure of T5 tail tip before and after interaction with FhuA in nanodisc. **A**. Negative stain EM image of phage T5, with the tail tip circled. **B**. Scheme of T5 tail tip with the assignment of the different proteins and their copy number. **C**. Cryo-EM structure of T5 upper tail tip at 3.5 Å resolution. Left: Isosurface view of the map seen from the side; middle: central slice side view; right: ribbon representation of the modelled proteins. The twist and rise between each ring is noted. **D**. Cryo-EM structure of the central fibre at 4.2 Å resolution. Left: isosurface view of the map seen from the side; right: ribbon representation of the modelled proteins. **E**. Map of T5 tail structural proteins genes. **F**. Negative stain EM image of a T5 tail interacting with a FhuA-nanodisc (black arrows). The white arrow points to the empty-filled limit of the tail (see also Extended Data Fig. 1F). The grey arrow head points to a density going through the nanodisc. **G**. Scheme of T5 tip after interaction with FhuA (Tip-FhuA). **H**. Cryo-EM structure of Tip-FhuA at resolutions ranging from 3.6 to 4.3 Å resolution. Isosurface view of a Tip-FhuA composite map seen from the side (left) and a central slice side view of it (middle). This composite map is formed by the addition of Tip-FhuA C3 open tube and C1 bent fibre maps, and is only for visualisation purposes; right: ribbon representation of the modelled proteins. The colour code in C, D, G, and H is the same as in B. Unattributed densities are in white. Bars: 50 nm.

## Results

### General architecture

T5 tails (11, 14) were preferred over whole phages for electron cryo-microscopy (cryo-EM) as they allow better quality imaging (Fig. S1A). Micrograph acquisition and extensive image processing (Fig. S2 and Table S1) yielded three maps of different tip subcomponents (Fig. 1C-D, Fig. S1C), whose resolution allowed to trace all the proteins from the tail tube to the distal end of the central fibre, except for STF_pb1_ and RBP_pb5_ (Table S1). After two TTP_pb6_ trimeric rings, the tube continues with a p140 trimer, then a DTP_pb9_ hexamer. A BHP_pb3_ trimer closes the tube and forms the beginning of the central fibre that continues with a pb4 trimer. The p140 trimeric ring is surrounded by a p132 dodecamer that forms the collar, onto which are grafted three trimeric STF_pb1_.

Upon T5 tail incubation with detergent-solubilised FhuA, BHP_pb3_ opens, TMP_pb2_ is expelled from the tube lumen and the central fibre disappears (14). As the presence of a lipid bilayer might stabilise a cell wall perforation intermediate, we used instead FhuA reconstituted into nanodiscs. Indeed, images of FhuA-nanodisc-incubated tails clearly show the presence of a nanodisc perpendicular to the tail tube at the rim of the open BHP_pb3_, a poorly defined protrusion going through the nanodisc, TMP_pb2_ partial ejection from the tail tube lumen and the bending of the central fibre with a very acute angle on one side of the tip (Fig. 1F, Fig. S1F). Extensive cryo-EM image processing yielded three other maps (Fig. 1H, Fig. S1H, S2, Table S1) allowing to trace all T5 tip proteins, except again STF_pb1_ and RBP_pb5_. Densities belonging to RBP_pb5_ are visible, but resolution is insufficient to build a model. However, a small angle neutron scattering envelop of the FhuA-RBP_pb5_ complex (19) could be very well fitted into the densities (Fig. S3D).

The density corresponding to the nanodisc is clearly visible (Fig. 2C,D), even though nanodiscs are heterogeneous in size and in position relative to the tail (Fig. S1F). Nanodisc density is not centred with respect to the tail tube axis: its centre of mass is shifted towards the bent fibre, below the density attributed to RBP_pb5,_ under which the structure of FhuA could be fitted (Fig. 2C-D, Fig. S3D). At low contour level, aligned with the tail tube lumen, a hole in the nanodisc is observed (Fig. 2D), strongly suggesting the presence of a channel at this position. At higher contour level, the tail tube lumen and the nanodisc are filled, and protrusions are visible above and below the nanodisc (Fig. 2C), as if a channel had perforated it. This channel is however poorly resolved, probably because of high heterogeneity in that region (Fig. S1F).

**Figure 2:**
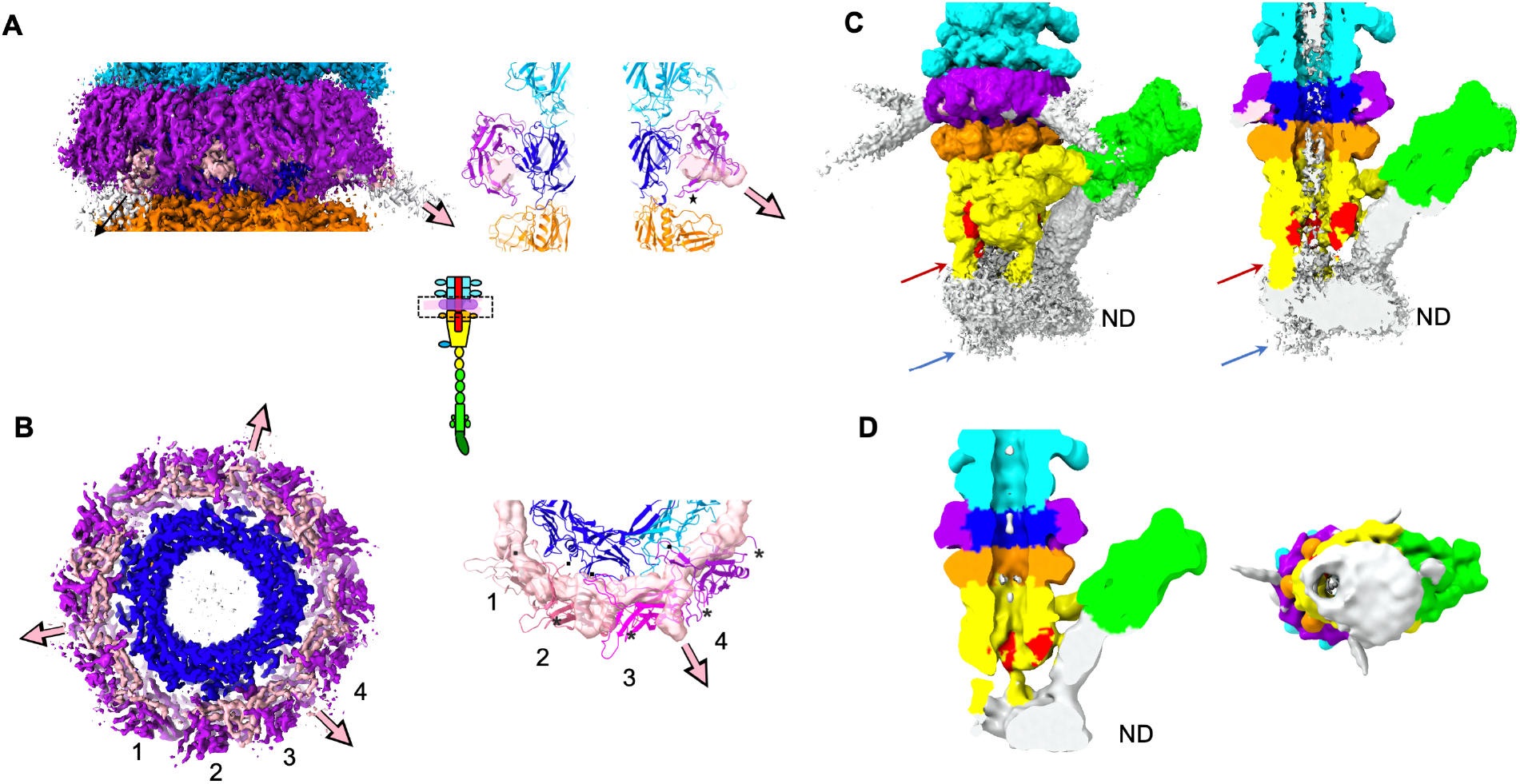
**A**. Left: Isosurface side view of the tip common core cryo-EM map at high contour level, centred on the p132 collar (boxed in the inset scheme of the tip). Right: ribbon representation of a central slice of the collar. The star points to loop 52-60 and the density attributed to STF_pb1_ is in transparent isosurface representation. **B**. Left: Isosurface bottom view of the previous map, slice at the p132 collar level. Right: bottom view of the four p132 monomers that are not related by the C3 symmetry (coloured from light pink to violet and numbered). They interact with two p140 monomers (cyan and blue) and with STF_pb1_ (transparent densities). The pink/black arrows point to the direction of the STF and the N- and C-termini of the proteins are respectively indicated by black dots and asterisks. **C**. Isosurface view at high contour level of Tip-FhuA unmasked and unfiltered cryo-EM map, side view (left) and slice (right). The red arrow points to one of the β-hairpin “leg”. The blue arrow point to the protrusion going through the nanodisc (ND). **D**. Isosurface view at a lower contour level of Tip-FhuA unmasked cryo-EM map, after a 15 Å lowpass filtering, slice (left) and view from beneath the nanodisc (right). The colour code is the same as in Fig. 1. Unattributed densities in C and D are in white.

Limited proteolysis experiments were performed on T5 tails and phage using subtilisin, and resulting particles analysed by SDS-PAGE and observed by negative staining EM. Within tail assembly, TMP_pb2_ is among the first to be digested (Fig. S4A,D), as it has a tendency to be expelled from the tail by its proximal extremity. This suggests that the protein is contained in the tube in a metastable state. On the contrary, proteins of the tail tip complex are extremely stable. TTP_pb6_ is also very stable with the exception of its decoration Ig-like domain. Upon analysis with negative stain EM, particles appear intact, with the exception of TMP_pb2_ in purified tails (Fig. S4B,E). Infectivity of proteolysed phages is also only mildly affected, decreasing by only an order of magnitude (Fig. S4C). Incubation with FhuA expels TMP_pb2_, making it even more susceptible to the subtilisin action, whereas elements of the tail tip remain resistant to proteolysis despite the vast conformational changes underwent.

### Core of the tip complex

The structures of the tip before and after interaction with FhuA share a common core, from the tail tube down to BHP_pb3_ and only start to diverge from BHP_pb3_ distal part. We previously determined a pseudo-atomic resolution structure of T5 tail tube: it exhibits a pseudo sixfold symmetry, the TTP_pb6_ gene resulting in a duplication/fusion of the canonical tail tube domains (TTD) gene (14) (Table S2). The TTD is formed by a β-sandwich flanked by an α-helix and a long loop (14)(Fig. S5A). At TTP_pb6_ C-terminus, an Ig-like domain decorates the tube, as in many Siphoviridae TTPs. Our tail tip structure includes two TTP_pb6_ rings that could be modelled (Fig. 1C). This higher resolution data on the TTDs confirms our previous modelling of the inter-ring interactions, mostly mediated by the long loops, the N-terminus, the linker between the two TTD and loops of the β-sandwiches (14), and have complementary charge surfaces (Fig. S6B). The rmsd between TTP_pb6_ structures of the proximal and distal ring is only 0.5 Å over all 464 residues (Fig. 5A) suggesting that the interface with the next p140 ring is very similar to that of two TTP_pb6_ rings. The densities of the Ig-like domains are poorly defined (Fig. 1C), witnessing a flexibility of this domain with respect to the tube scaffold (23). The tube extends through the tip: after the last TTP_pb6_ ring, it continues with a pseudo-hexameric p140 ring, a hexameric DTP_pb9_ ring and the proximal domains of the BHP_pb3_ trimer forms the last pseudo-hexameric ring of the tube. The structure of these proteins is also based on the canonical TTD (6) but differently decorated. Thus, the tube diameter is conserved, even though the pitch and the twist between the different rings are different (Fig. 1C). As for TTP_pb6_, the interaction between the rings is mediated mostly by the long loops, the N-terminus and loops of the β-sandwiches (14); they also have complementary charged surfaces (Fig. S6B). The inner surface of the tube is highly electronegative until DTP_pb9_ (Fig. S6A) allowing DNA to slide along it (6).

p140 pairwise comparison with TTP_pb6_ results in a very high DALI (24) Z-score (Fig. S5B), pointing to TTP_pb6_ gene duplication to form p140, despite identity between the two proteins being only 9% (Fig. S5C). The main difference between the two proteins is the absence of the Ig-like domain in p140. Indeed, the p140 ring is surrounded by a larger p132 dodecameric ring, p140 C-terminus making direct contacts with a p132 monomer (Fig. 1C, 2A,B), explaining the need for a decoration-less ring at this position. As suggested by p140 gene position in T5 genome (Fig. 1E), this protein is a *bona fide* component of its baseplate. p140 and p132 genes are a landmark of the large T5-like phages family only. It thus seems that STF anchoring could occur differently in other *Siphoviridae*, in particular in the lambdoid phages. In the *Myoviridae* phage T4, the presence of an additional ring of the TTP-like protein gp54 between the ‘ring initiator’ DTP_gp48_ and the first *bona fide* TTP_gp19_ rings (25) is also observed, and a role in sheath assembly initiation has been proposed (31). This additional ring is also not systematically present within the Myoviridae family.

We previously determined the crystal structure of DTP_pb9_, showing that the DTP was a common feature to both Gram-positive and Gram-negative infecting siphophages (17). In all other phages and tail-like machines, this protein ensures the six-to-threefold symmetry transition between the TTP hexamer and the BHP trimer. Here however, the DTP_pb9_ ring is sandwiched between two threefold symmetric rings, which both have a pseudo-sixfold symmetry. This explains the low rmsd between the two DTP_pb9_ monomers that are not related by the imposed threefold symmetry of the map. DTP_pb9_ TTD is decorated with an OB (Oligonucleotide-Oligosaccharide Binding) domain (Fig. S5D). The OB domains, as the Ig-like domains, are proposed to interact with carbohydrates at the cell surface and serve to increase infectivity (4). Unlike DTPs of siphophages infecting Gram-positive bacteria that serve as a platform to anchor the RBPs (26–28), DTP_pb9_ does not bind any other protein than those forming the tail tube.

### The collar, p132 and STF_pb1_

The collar is made of a p132 dodecamer. The p132 fold belongs to the Immunoglobulin superfamily (Fig. S5E) and a DALI search links it to the N-terminal domain of the Baseplate Protein Upper (BppU, ORF48) of phage TP901-1 (27). Ig-like domains in phages are usually decoration domains, here however, as for BppU, it is a bona fide structural protein that serves to anchor STF_pb1_. The dodecameric p132 ring completely surrounds the trimeric p140 ring, with p132-p132 and p132-p140 contacts in both p132 and p140 proximal regions and, to a much lesser extent, p132-TTP_pb6_ contacts, mainly mediated through the loops and termini of the three proteins (Fig. 1C, 2A,B). There are no interactions between p132 and DTP_pb9_ or the Ig-like domain of TTP_pb6_, as determined by PISA (29).

Unattributed densities in the lower part of the collar, intertwined between the p132 monomers, most probably belong to STF_pb1_. However, map quality/connectivity did not allow to unambiguously model it, but it could correspond to its ∼50 N-terminal residues. These densities point out of the collar to form the start of the three STF. The intricate STF_pb1_-p132 interaction (Fig. 2A,B) explains that in the fibreless T5-hd1 mutant, which bears a mutation in pb1 gene leading to a truncated protein, the collar is absent and the p132 protein is not detected by Western blotting. Also, p132 could not be localised *in phago* using anti-p132 IgGs (11). In solution, p132 bears a folded core with very flexible loops and termini (30), which could be the target of the rabbit IgGs. These loops are mainly unavailable *in phago*, as they are involved in protein-protein interactions (Fig. 2B).

Four consecutive p132 monomers, not related by the threefold symmetry, show high structural similarity (Fig. 2B, Fig. S5E,F), even though their environment is different. Indeed, there is a symmetry mismatch between STF_pb1_ threefold, p140/TTP_pb6_ pseudo-sixfold and p132 pseudo-twelvefold symmetries. This symmetry mismatch is absorbed by the loops and N-termini of p132 monomers. Finally, the C-terminus and loop 52-60, involved in p132-p132 interactions are less variable (Fig. S5E).

### Closing the tube: BHP_pb3_ and TMP_pb2_

BHP_pb3_ trimer forms the most distal TTD ring of the tube (Hub Domains (HD) I and IV, Fig. 3A, Table S2). The linker between the two TTDs and the long loop of the second TTD have evolved into larger domains (HDII and insertion, and HDIII respectively, Fig. S7A,B), the first of which is large enough to close the tube *via* a plug domain (Fig. 3A). A long linker runs along the protein down to the tip of the cone, inserted between two neighbouring BHP_pb3_ subunits and contributing to the stability of the closed tube (Fig. 3A). BHP_pb3_ C-terminus forms the beginning of the central fibre with two fibronectin domains (FNIII), further stabilising the closed tube (Fig. 3B) and giving BHP_pb3_ the shape of a trophy cup (Fig. 1C,D, 3A). BHP_pb3_ results from a TTD duplication-fusion, as TTP_pb6_. These two duplication/fusion events are clearly independent however, as the fusion did not occur in the same way in the two proteins (Fig. S7B).

**Figure 3:**
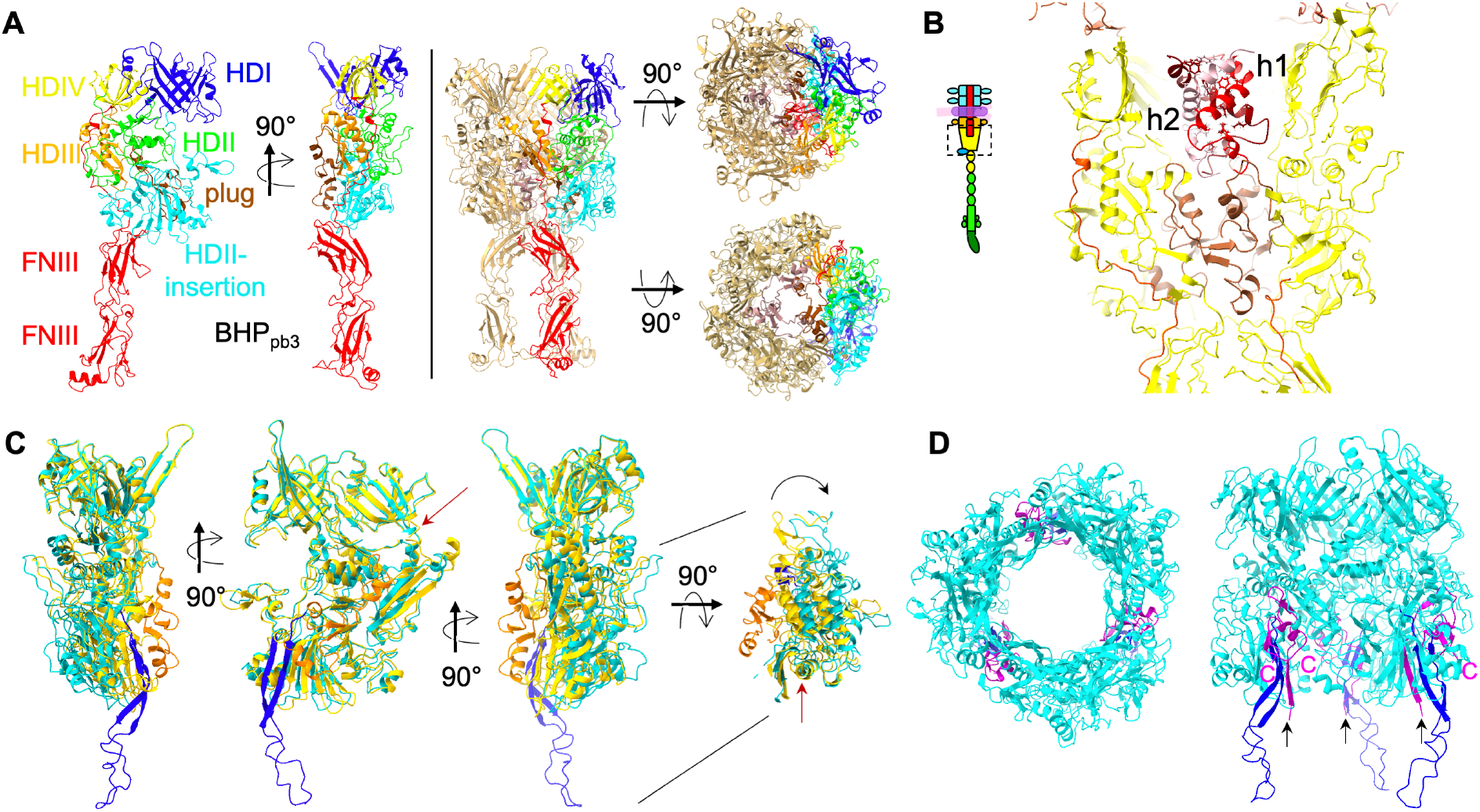
BHP_pb3_ closing and opening of the tube. **A**. Left: Two side views of a BHP_pb3_ monomer in ribbon representation, with HDI-IV domains coloured blue, green, orange and yellow, respectively, the HDII insertion in cyan, the plug in brown and the C-terminus extension, comprising the HDIV-FNIII linker and the two FNIII, in red. Right: side, top and bottom views of the BHP_pb3_ trimer. One monomer is coloured as on the left panel and the three plug domains are coloured brown. In the bottom view, the FNIII have been removed for clarity. **B**. Central slice through BHP_pb3_ cup, boxed in the inset scheme of the tip (yellow, HDIV-FNIII linker: red, plug: brown) highlighting the 35 resolved residues of TMP_pb2C_ in different shades of red. Hydrophobic residues of TMP_pb2C_, pointing to the centre of the coil are represented in sticks. **C**. Overlay of BHP_pb3_ before (yellow, plug: orange) and after (cyan, plug: blue) opening of the BHP_pb3_ cone, after superimposition of the tip. Three side views 90° apart and a top view are shown. In the top view, HDI and HDIV have been removed to highlight the pivotal movement of the HDII-insertion domain. A red arrow points to the long helix of HDIII that acts as a hinge (see also Movie S4). The long linker and the FNIII have been removed for clarity. **D**. Top and side views of the open BHP_pb3_ trimer with the same colour code as in C, and TMP_pb2*_ 42 C-terminal residues in magenta. TMP_pb2*_ C-termini are indicated (C) as well as the last built residue in N-terminal (T1085, black arrow).

Unexpectedly, on unsymmetrised EM reconstructions, an extra density is observed at the base of only one BHP_pb3_ monomer (Fig. S3A). Resolution was insufficient to build an atomic model de novo, but secondary structure features could be identified. From the eleven proteins that form the tail and that have been identified by mass spectrometry (MS) (Table S3), only p143 has not been located. Its gene position in the tail structural module (Fig. 1E) suggests it is the Tail Completion Protein, located in the head-to-tail joining region (4, 6). However, we have solved T5 proximal tail region and could not identify it there (Linares et al, in preparation). A flexible fit of an Alphafold2 (31) structure prediction of p143 into this extra density was convincing (Fig. S3B). Thus, we propose that this unattributed density corresponds to p143. This density is absent from reconstructions of the tip after interaction with FhuA, suggesting that the protein could detach during the infection process.

The tube lumen is filled with TMP_pb2_. TMP_pb2_ (15, 32), as λ TMP (33), undergoes proteolytic cleavage during tail morphogenesis, but it was supposed that the cleaved peptide, TMP_pb2C_, was removed from the final tail assembly, leaving the rest of the protein, TMP_pb2*_, in the tail tube lumen. TMP_pb2_ density is very ill-defined along the tail tube, probably due to a poor interaction network between the predicted coiled-coil (15) of TMP_pb2_ and the wall of the tube, except in the BHP_pb3_ cup (Fig. 1C, 3B). There, we could model TMP_pb2C_ 35 C-terminal residues. Three TMP_pb2C_ copies are coiled in a superhelix, burying hydrophobic residues in its centre, interacting closely with BHP_pb3_ plug (Fig. 3B). TMP_pb2C_ presence in the tail is confirmed by proteomics and LC-ESI-TOF-MS (Table S3, Fig. S8C), and allowed us to localise TMP_pb2_ cleavage site (Fig. S5G). This latter is located between the TMP_pb2*_ hydrophobic stretch and a metallopeptidase motif that was shown to have muralytic activity (15), separating this enzymatic domain from the rest of the protein. This is reminiscent of the situation in T4, where the cell puncturing protein gp5 is cleaved during tail assembly between its lysozyme domain and the β-helix spike (34). It was previously suggested that TMP_pb2_ (32), as other phages TMPs (e.g. λ (35)) is a hexamer, but our data clearly indicates that only a trimer is present. A trimer of the 20 C-terminal residues of phage 80α TMP was also modelled (26), suggesting that this might be a general feature of siphophages. Interactions between TMP_pb2_ and BHP_pb3_ are mainly electrostatic (Fig. S6B,C), unlike in 80α.

### The central fibre and its re-arrangement upon receptor binding

BHP_pb3_ C-terminal ∼210 residues and pb4 N-terminal ∼350 residues were modelled *de novo* (Fig. 1C,D). As the resolution of the fibre map drops towards the fibre tip because of its flexibility, the rest of pb4 protein was modelled using flexible fitting of the better resolved equivalent domains built in the Tip-FhuA maps (see below and Fig. S1H).

The proximal region of the central fibre is made of three strings of five consecutive FNIIIs, two at BHP_pb3_ C-terminus and three at pb4 N-terminus. It starts as three independent strings; the repulsion between them could be caused by an important negative patch at the surface of the BHP_pb3_ FNIIIs (Fig. S6D). BHP_pb3_-pb4 interaction is ensured by two distal loops of the second FNIII of BHP_pb3_, and the N-terminus and two proximal loops of the first pb4 FNIII (Fig. S8A). After a hinge region, the three pb4 monomers merge to form a 110-Å long β -helix spike, formed of a 24 β-strand longitudinal mixed β-sheet prism. It has a triangular section with a mean diameter of 20 Å, delineating a very dense and hydrophobic interior (Fig. 4B). β-helix spikes/fibres are very common in phage host-recognition or/and perforation apparatus and a DALI search indeed relates pb4 spike to different phage and tail-like machines spikes/fibres (Fig. 8B). Here however, pb4 does not have a role in perforation or recognition.

**Figure 4:**
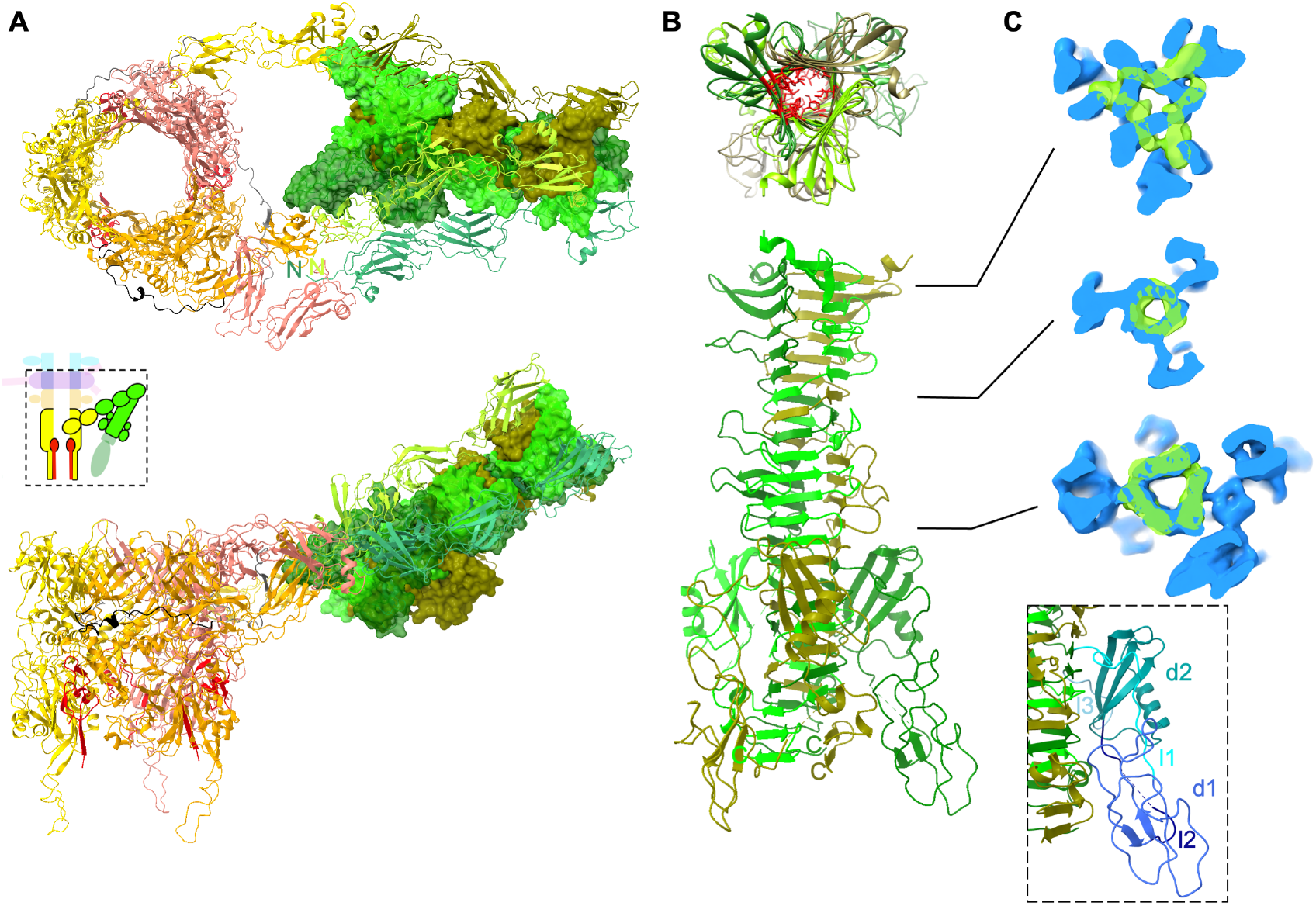
Bending of T5 straight fibre. **A**. Structure of Tip-FhuA BHP_pb3_, 42 C-terminal TMP_pb2*_ residues and pb4 (boxed in the inset scheme of the tip). BHP_pb3_ monomers are in gold, orange and salmon with the HDIV-FNIII linker coloured in different shades of grey, pb4 in different shades of green and TMP_pb2*_ in red. All proteins are in ribbon representation except for pb4 spike that is in surface representation. Top: top view. pb4 N-termini (N) and BHP_pb3_ C-termini (C) are indicated; bottom: side view. **B**. Top: top view of pb4 spike. pb4 monomers are in different shades of green. The hydrophobic residues pointing towards the interior of the spike are depicted red and in sticks in one subunit only. Bottom: Side view of pb4 spike. C-termini are indicated (C). **C**. Superimposition of pb4 spike in isosurface view of the tip (green) and Tip-FhuA (blue) maps (unsharpened). Three slices are shown and their position along the spike is indicated in b. The map after interaction with the receptor contains the spike decoration domains and the FNIIIs while that before interaction contains only pb4 spike. Inset: pb4 spike decoration domains and linkers are coloured in different shades of blue on one pb4 subunit (linker 1 (l1, residues 484-547), domain 1 (d1, 548-566), linker 2 (l2, 548-566), domain 2 (d2, 567-618) and linker 3 (l3, 519-626)). A DALI search links domain 2 to phage spike decoration domains (PDB codes 7CHU-A, 6TGF-D, 6E1R-A, 5M9F-A, 6NW9-C, 6EU4-A, 5W6H-A with DALI Z-score of 3.9 to 2.3, rmsd from 2.4 to 3.0 Å over ∼45 residues and identity ranging from 4 to 14%).

The spike is decorated with two small globular domains inserted between β-strands 15 and 16 of each pb4 subunit, and which are connected by relatively long linkers (Fig. 4B). These domains are not visible in the central fibre map, probably because of the degrees of freedom offered by the linkers. A DALI search links the second domain to decoration domains found in other tail spikes.

The central fibre ends with RBP_pb5_ but this part of the map is poorly resolved and did not allow to build an atomic model of this protein. This 310 Å-long central fibre bears two hinges, one between pb4 last FNIII and the spike and the other at the spike-RBP_pb5_ interface: it introduces some controlled flexibility to this otherwise rigid assembly and may ease RBP_pb5_ exploring space and encountering its bacterial receptor.

Upon RBP_pb5_ binding to FhuA, the central fibre reorganises: the three FNIIIs strings dissociate, two strings relocate on one side of BHP_pb3_ and the third one on the other side (Fig. 4A, Movies S1, S2). This reorganisation of the central fibre is allowed by the long linker that connects BHP_pb3_ HDIV to the first FNIII, and which now runs perpendicular to the tail axis along BHP_pb3_. The central fibre bends by ∼160° at the level of the hinge between the third pb4 FNIII and pb4 spike: the latter is now surrounded and stabilised by pb4 FNIIIs, which interact with and stabilise pb4 spike decoration domains (Fig. 4A). pb4 spike also undergoes structural rearrangement, with a different twist of the spike (Fig. 4C). This bending and stabilisation of the central fibre bring the tail tube closer to the membrane, orient it correctly and anchor the tail to the membrane.

### Tube opening and anchoring of the tail to the membrane

As mentioned above, the structures of the tip before and after interaction with FhuA start to diverge from BHP_pb3_ distal domains. More precisely, HDI and HDIV overlay remarkably (Fig. 3C, Movie S3), with a rmsd of 0.85 Å over 244 residues. However, HDII-III rotate around the long helix of HDIII, as a rigid body (rmsd between HDII-III before and after opening is 0.75 Å over 367 residues), and the plug unwinds in a long β-hairpin (Fig. 3C,D, 4a and Movies S3, S4). These conformational changes result in BHP_pb3_ trimer opening, creating a channel with a constant ∼40 Å diameter from the HDI-HDIV ring to the HDII-insertion tip (Fig. 3C,D, Movie S4). The three β -hairpins ‘legs’ connect BHP_pb3_ to the nanodisc (Fig. 2C,D): indeed, their tip is composed of 233-Leu-Phe-Gly-Leu-236, which would insert into the outer leaflet of the lipid bilayer hydrophobic core. Above these hydrophobic residues stand 230-Lys-Lys-Lys-232 and Arg238, conferring a strong positive charge to the β-hairpin (Fig. S6E). They could interact with the negatively-charged phosphate groups of the lipopolysaccharides, further stabilising the anchoring of the β-hairpin to the membrane.

In the crevice opened at the interface between two BHP_pb3_ subunits, extra densities were identified, in which the 43 C-terminal residues of TMP_pb2*_ could be modelled (Fig. 3D, 4A). These densities merge with the ill-defined densities of BHP_pb3_ β-hairpin, strongly suggesting that TMP_pb2*_ continues towards the nanodisc along BHP_pb3_ β -hairpin, forming with the latter a 3-stranded β -sheet. Indeed, TMP_pb2*_ continues with a stretch of 9 residues, long enough to reach the nanodisc, followed by a stretch of 46 hydrophobic residues, compatible with two transmembrane helices (Fig. S5G), which would insert into the outer-membrane and form a channel (Fig. S3C).

Thus, T5 tail tube is anchored to the outer-membrane by both BHP_pb3_ and TMP_pb2*_, in addition to FhuA-RBP_pb5_: it ensures that the tail tube is locked in register with the channel formed by TMP_pb2*_ in the outer-membrane.

## Discussion

### Baseplate comparison

As expected, BHP_pb3_ structure partially aligns with other phages and tail-like machines BHPs, with high DALI Z-scores (Fig. 7C): the four HD of the canonical T4 BHP_gp27_ (*34*) are also present in BHP_pb3_, but there is a large insertion in HDII, which closes the tube (Fig. 3A, Fig. S7D). In *Myoviridae* and tail-like machines, the tail tube is closed by an OB domain followed by a spike (7–9, 34) (Fig. S7D). In *Siphoviridae*, there is more diversity for closing the tail tube. Indeed, the four baseplate structures available to date (phages T5, 80α (26), p2 (28), and GTA (36)) exhibit three different closing modes (Fig. 7D): in p2 BHP, two HDII loops pointing towards the lumen of the tube are longer than in *Myoviridae* and close the tube. There, tube opening is induced by an iris-like movement, triggered by Ca^2+^ binding, of HDII-III (28). In 80α, the tube is closed by a helix in the C-terminal extension of the BHP that forms a twisted tripod in the trimer lumen (26). Finally, T5 and GTA tubes are closed by the large HDII insertion (36). Indeed, the two proteins composing GTA BHP, the Hub and the Megatron, align extremely well with BHP_pb3_ (Fig. S7C,E).

Superimposing the *Siphoviridae*-related baseplate structures available, it was striking to observe the remarkable structural superposition of helix 2 of TMP_pb2C_, the resolved helix of TMP_80α_ and helix α1 of the Iris/penetration domain of GTA Megatron (Fig. S5H). Sequence alignment showed no detectable sequence similarity, and in the case of T5 TMP_pb2C_ and TMP_80α_ C-terminus, the interaction with the BHP is *via* the bottom of the BHP cup (Fig. 1C, 3B). In GTA, helix α1 of the Megatron is proposed to be a pore-lining helix that could insert in the outer-membrane to allow the DNA across it. This however cannot be a general feature in *Siphoviridae*.

### T5 trigger for infection and formation of a channel

Our data allow to propose a mechanism for T5 trigger for infection and formation of a channel (Fig. 5). Upon RBP_pb5_ binding to FhuA, a conformational change at the RBP_pb5_-pb4 interface would occur, resulting in a different twist of pb4 spike in its proximal part. This twisting of the spike would pull on pb4 FNIII-spike linker, leading to the disruption of the FNIII-spike interaction network (Fig. 5B). The association between this FNIII and the spike is thus loosened, the pb4 FNIII-spike linker reorganises and stabilises a new interaction network between the FNIIIs and the spike (Fig. 5C). This series of events results in the bending of the central fibre, at the level of the FNIII-spike hinge, on the side of the tube, pulling the open tube towards the host membrane (Fig. 5D).

**Figure 5:**
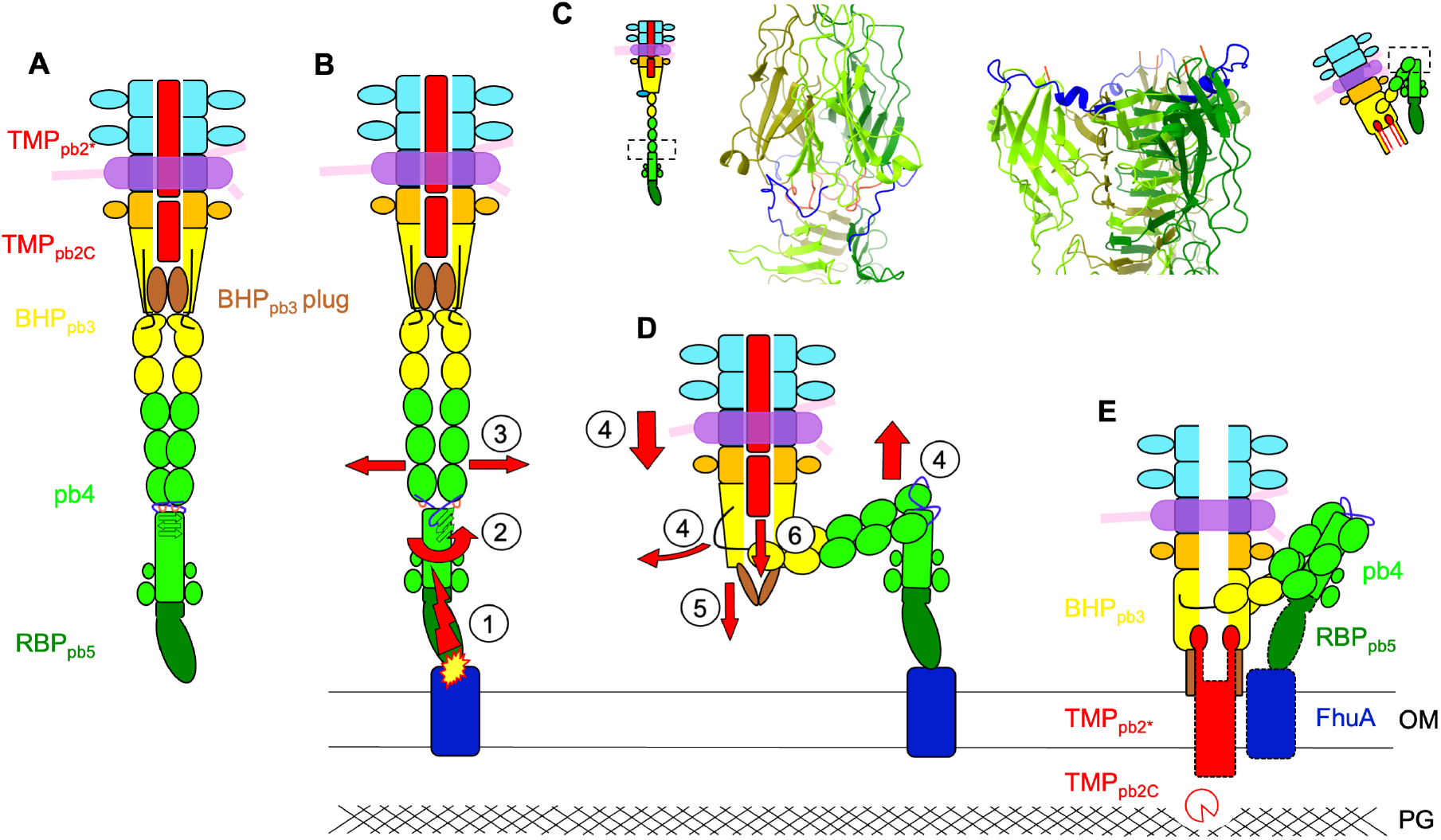
Proposed mechanism of trigger for infection. **A**. Scheme of T5 tail tip. The HDIV-FNIII linker (black) and the plug (brown) are highlighted in BHP_pb3_, and the FNIII-spike linker (blue), loop 224-232 of the third FNIII (salmon) and the orientation of the proximal three β-strands of the spike (black arrows) in pb4. **B**. Following RBP_pb5_-FhuA interaction, a conformational change (1) would induce a twisting of the proximal pb4 spike (2), pulling on pb4 FNIII-spike linker. This in turn would destabilise the FNIII string network (3). **C**. Blow up on pb4 FNIII-spike interface before (left) and after (right) interaction with FhuA. The two spikes are aligned on the middle sheet of the spike (residues 413-465). pb4 subunits are in different shades of green, the FNIII-spike linker in blue and FNIII loop 224-232 in salmon. **D**. The FNIII string reorganisation around pb4 spike would induce pb4 bending, bring the tube close to the membrane and disengage BHP_pb3_ HDIV-FNIII linker (4). This latter event would liberate the plug, opening the tube (5) and destabilising TMP_pb2C_, which would be expelled (6). **E**. BHP_pb3_ plugs refold as a β-hairpin legs and anchor in the outer-membrane (OM), TMP_pb2*_ is also expelled, its C-termini inserting in the crevice between BHP_pb3_ subunits, its hydrophobic segment inserting in the OM to form a channel. TMP_pb2C_, released in the periplasm, would digest the peptidoglycan (PG). In **E**, coloured boxes depict proteins that could be modelled (full line) or for which densities are visible (dotted line). TMP_pb2C_, for which no densities is visible but for which we propose a location, is represented as an empty Pacman.

To accommodate for the drastic conformational changes of the FNIII strings induced by the bending of the central fibre, BHP_pb3_ HDIV-FNIII linker is pulled and, like a zip, disrupts the interaction network between BHP_pb3_ monomers and with BHP_pb3_ plug. This then allows the rotation of BHP_pb3_ HDII-III, plug unfolding/refolding, opening of the tube and anchoring of BHP_pb3_ to the membrane *via* the β-hairpins legs (Fig. 5D). BHP_pb3_ closed conformation would be stabilised in a metastable state by the assembly process and interaction with its tip partners, TMP_pb2C_ in particular. Open BHP_pb3_ would be of lower energy and would drive the conformational changes underwent by the tip complex upon infection. T5 tip structure was proposed to the CASP14 competition: interestingly, BHP_pb3_ *open* structure only was correctly predicted (37).

BHP_pb3_ opening disrupts the interaction with TMP_pb2C_, which would be expelled from the tube and translocated to the host periplasm where it would locally digest the peptidoglycan; a refolding step could be necessary. TMP_pb2*_ would in turn be expelled from the tail tube and anchor its C-terminus in the crevice created between BHP_pb3_ monomers upon opening (Fig. 5E). Its hydrophobic segments would then insert in the outer-membrane to form a transmembrane channel. These events would be thermodynamically favoured by hydrophobic segments insertion in the membrane and TMP_pb2*_ alleged metastable state within the tail tube. TMP_pb2_ was shown to be involved in contact points between outer- and inner-membranes (38). Whether TMP_pb2*_ inserts into both the outer- and the inner-membrane remains to be determined: to form a channel wide enough to allow DNA through would require the six TMP_pb2*_ transmembrane helices. Insertion into the inner-membrane could thus occur *via* another part of the protein and/or the recruitment of host proteins.

The mechanism presented in this paper, by which receptor binding triggers the opening of its tail tube, its anchoring to the host membrane and formation of a transmembrane channel is, to our knowledge, the first one described for *Siphoviridae*, the most prevalent family of phages. It is furthermore entirely original compared to what was known and described so far for the more complex *Myoviridae* and related bacterial machines. Our study provides a solid structural basis to further explore the diversity of viral entry mechanisms and their properties.

## Acknowledgments

We acknowledge the European Synchrotron Radiation Facility for provision of beam time on CM01. This work used the platforms of the Grenoble Instruct-ERIC center (ISBG; UMS 3518 CNRS-CEA-UGA-EMBL) within the Grenoble Partnership for Structural Biology (PSB), supported by FRISBI and GRAL, financed within the University Grenoble Alpes graduate school (Ecoles Universitaires de Recherche) CBH-EUR-GS. The IBS acknowledges integration into the Interdisciplinary Research Institute of Grenoble (IRIG, CEA). We thank Luca Signor, Emmanuelle Neumann and Daphna Fenel for technical assistance, Leandro Estrozi, Ambroise Desfosses, Benoît Arragain for help and discussion on image analysis, Aymeric Peuch for help with the EM computing cluster, Christophe Masselon, Kavya Clement and the EdyP MS platform for MS analysis and Christine Ebel for useful discussions.

## Author Contributions

CB conceived the project. CAA, RL, CD and CB prepared T5 tails and FhuA-nanodiscs used for high-resolution cryo-EM. GS and RL optimised cryo-grid preparation. GE recorded the cryo-EM data. GE, CAA and RL processed the cryo-EM data. RL built the atomic models. CB analysed and interpreted the models with the help of CAA. CD, CAA and RL performed the limited proteolysis experiments. NHE and EBE performed and analysed the MS data. CB wrote the paper, with contributions from RL, CAA and GS. All authors contributed to the editing of the manuscript.

## Data and materials availability

Cryo-EM density maps of T5 tip resolved in this study and the associated atomic coordinates have been respectively deposited in the Electron Microscopy Data Bank (EMDB) and in the Protein Data Bank (PDB) under the following accession codes: EMD-13953 / PDB 7QG9 (Tip / Tip-FhuA common core), EMD-14733 / PDB 7ZHJ (Tip without fibre), EMD-14790 / PDB 7ZLV (Tip fibre), EMD-14869 / PDB 7ZQB (Tip full), EMD-14799 / PDB 7ZN2 (Tip-FhuA full), EMD-14800 / PDB 7ZN4 (Tip-FhuA bent fibre) and EMD-14873 / PDB 7ZQP (Tip-FhuA open tube). See also Table S1.

## Materials and Methods

### T5 tail purification

T5 tails were preferred over whole phages for cryo-EM as the former allow thinner ice and no DNA background, and thus better quality images. *E. coli* strain F cultures at 37°C were infected during the exponential growth phase with the amber mutant phage T5D20*am*30d, which bears an Amber mutation in the major capsid protein gene, at a multiplicity of infection of 6-7. After complete cell lysis (OD600nm < 0.15), the cell lysate was incubated with a pinch of DNAse and 5 mM MgCl_2_ at 37°C for an hour and centrifuged 20 min at 6000 rpm to remove cell debris and unlysed cells. T5 tails were then precipitated from the culture medium by incubation with 0.5 M NaCl and 10 % (w/w) PEG 6000 overnight at 4°C. The pellet of a 20 min 8000 rpm centrifugation was resolubilised in 10 mM Tris pH 8, 100 mM NaCl 1 mM CaCl_2_, 1 mM MgCl_2_ and purified on a glycerol step gradient (10 to 40 %) in the same buffer centrifuged 2 h at 20,000 rpm (SW41 rotor). The gradient fractions containing the tails (usually ∼10 % glycerol), diluted ten times in 10 mM Tris pH 8, were loaded onto an ion exchange column (HiTrap™ Q HP 1 mL, GE Healthcare) equilibrated and washed in the same buffer. The tails were eluted by a 0 – 0.5 M NaCl linear gradient. Purified tails were incubated 30 min with FhuA loaded nanodiscs in a tail:FhuA-nanodisc ratio of 1:10 (vol/vol) at room temperature. The produced tails were shown to behave as capsid attached tails: in particular, they can interact and perforate outer-membrane vesicles (11) and interaction with the purified, detergent-solubilised receptor FhuA results in the expulsion of the TMP_pb2_ and the loss of the central fibre (14).

### FhuA containing nanodiscs

The gene coding for the Membrane Scaffold Protein MSP1E3D1 (*39*) was cloned in a pET28a plasmid, used to transform BL21(DE3) *E. coli*. Protein expression was induced by the addition of 1 mM IPTG to a 37°C-growing TB/kanamycin (50 μg.mL^-1^) culture when it reached an OD_600nm_ of 1.2. Cell were harvested 4 h later, resuspended in a lysis buffer (20 mM NaPO4 pH 7.4, 1 % (w/v) Triton X-100, lysozyme 0.2 mg.mL^-1^, DNase I 0.2 mg.mL^-1^) and broken through 3-4 passages in a microfluidizer (13 kpsi). After clarification of the cell lysate, the supernatant was loaded on a nickel affinity column (HiTrap™ Chelating HP 5 mL, GE Healthcare) previously equilibrated with 40 mM Tris-HCl pH 8.0, 300 mM NaCl, 1% (w/v) Triton X-100. The column was then washed with the same buffer, then with 40 mM Tris-HCl, pH 8.0, 300 mM NaCl, 50 mM Sodium Cholate, then 40 mM Tris-HCl pH 8.0, 300 mM NaCl and then 40 mM Tris-HCl pH 8.0, 300 mM NaCl, 10 mM Imidazol. The protein was eluted with 40 mM Tris-HCl, pH 8.0, 300 mM NaCl, 300 mM Imidazol. The histidine tag was cleaved by incubating overnight the MSP with the TEV protease in a MSP:TSV 1:10 (w/w) ratio at room temperature in 50 mM Tris-HCl, pH 8.0, 0.5 mM EDTA, 1 mM DTT and then dialysed 2 h against 20 mM Tris-HCl, pH 8.0, 150 mM NaCl. The cleaved protein was loaded onto the same nickel affinity column equilibrated with 20 mM Tris-HCl, pH 8.0, 150 mM NaCl. Cleaved MSP was recovered in the flow through, concentrated and loaded onto a size exclusion column (SD75 10/300 GL, GE Healthcare) equilibrated in 20 mM Tris-HCl, pH 7.4, 150 mM NaCl, 0.5 mM EDTA. Fractions of pure protein were pooled and the protein was concentrated to 2 mg/ml on Amicon concentrator (MWCO 100 kDa).

FhuA was produced and purified as described (18) : *E. coli* AW740 {FhuA31 ΔompF *zcb*::Tn*10* ΔompC} transformed with the pHX405 plasmid, in which the *fhuA*, gene under control of its natural promoter, was grown at 37°C in LB medium supplemented with ampicilin (125 μg.mL^-1^), tetracyclin (10 μg.mL^-1^) and 2,2’ bipyridyl (100 μM), an iron chelator used to induce FhuA production. After clarification of the cell lysate, total membranes were recovered by ultracentrifugation and solubilised using 50 mM Tris pH 8.0, 2% (w/w) OPOE (N-octylpolyoxyethylene, Bachem) at 37°C for half an hour. The insoluble material was recovered by ultracentrifugation and solubilised 1 h at 37°C using 50 mM Tris pH 8.0, 1 mM EDTA, 1% (w/w) LDAO (N,N dimethyl dodecylamine-N-oxide, Anatrace). The solubilised fraction, recovered after ultracentrifugation, was supplemented with 4 mM MgCl_2_ and 5 mM imidazole and loaded on a nickel affinity column (HiTrap™ Chelating HP 5 mL, GE Healthcare) previously equilibrated with 0.1 % LDAO, 20 mM Tris pH 8.0, 200 mM NaCl and washed with the same buffer. The protein was eluted from the column with 0.1% LDAO, 20 mM Tris pH 8.0, 200 mM imidazole, and loaded onto an ion exchange column (HiTrap™ Q HP 1 mL, GE Healthcare) equilibrated with 0.05% LDAO, 20 mM Tris pH 8.0. The protein was eluted by a 0 – 1 M NaCl linear gradient.

To produce FhuA loaded nanodiscs, purified FhuA was incubated with DOPC (1,2-dioleoyl-sn-glycero-3-phosphocholine) solubilised in 100 mM Sodium Cholate and MSP1E3D1 in a 1:6:360 FhuA:MSP:lipid molar ratio. After 1 h incubation at 4°C, detergent was removed by the addition of 50 mg.mL^-1^ BioBeads (BioRad) and incubation on a stirring wheel at room temperature for 2 h. FhuA loaded nanodiscs were further purified on a nickel affinity column equilibrated in 20 mM Tris pH 7.5, 150 mM NaCl, 5 mM Imidazol, eluted with the same buffer containing 200 mM Imidazol, and desalted on a PD10 (GE Healthcare) desalting column equilibrated in 10 mM Tris pH 7.5, 100 mM NaCl, 1 mM CaCl_2_, 1 mM MgCl_2_.

### Cryo-EM sample preparation

Typically, 3.5 μL of T5 tail sample (with or without FhuA-nanodisc) were deposited on a freshly glow discharged (25 mA, 30 sec) Cu/Rh 300 mesh Quantifoil R 2/1 EM grids and plunge-frozen in nitrogen-cooled liquid ethane using a ThermoFisher Mark IV Vitrobot device (100% humidity, 20°C, 5 s blotting time, blot force 0).

### EM data acquisition

Respectively 3208 and 9608 micrographs (splitted over two data collections for the latter) were collected for tails alone and tails incubated with FhuA-nanodisc. 40-frame movies were acquired on a Thermo Fisher Scientific Titan Krios G3 TEM (European Synchrotron Radiation Facility, Grenoble, France)(40) operated at 300 kV and equipped with a Gatan Quantum energy filter coupled to a Gatan K2 summit direct electron detector. Automated data collection was performed using Thermo Fisher Scientific EPU software, with a typical defocus range of -1.0 to -3.0 μm and a total dose of 40 e^-^/Å^2^ per movie. A nominal magnification of 105.000x was used, resulting in a calibrated pixel size at the specimen level of 1.351 Å.

### EM image processing

Frame alignment was performed using Motioncor2 (41) keeping respectively frames 3 to 30 and 1 to 40 for Tip and Tip-FhuA, and applying dose weighting. Contrast Transfer Function parameters were then determined using Gctf (42); Manual particle picking was performed with EMAN2/e2helixboxer (43). The first picking coordinate was consistently centred on T5 collar and the second one a few hundred Å toward BHP_pb3_, along the central fibre (Tip) or the tail axis (Tip-FhuA) (Extended Data Fig. 2). This “vectorial” picking allowed to choose and adapt the position of the box along that axis prior to extraction and proved to be very efficient. All subsequent image processing was performed using Relion (versions 3.0 and 3.1)(44). Flowchart of the EM processing pipeline is presented in Extended Data Fig. 2.

### Tip

After particle extraction (box size of 340x340 pixels^2^) centred 80 Å under the collar and 2D classification, a homogeneous dataset of 9,290 particles was obtained. No 3D classification was performed. Using a 15 Å resolution map determined from a previous cryo-EM data collection (45) as an initial model, a C3 reconstruction of the tip was calculated, from the 2^nd^ TTP_pb6_ ring to the beginning of the central fibre. After masking and sharpening, the overall estimated resolution of the map reached 3.53 Å (FSC_0.143_). A new set of particles (box size of 400x400 pixels^2^) was extracted after a 150 pixels coordinate shift on the z-axis, toward RBP_pb5_. A 15 Å low-pass filtered initial model was generated from the newly extracted particles using relion_reconstruct tool and determined a C3 reconstruction of the central fibre, from BHP_pb3_ to the beginning of RBP_pb5_. After masking and sharpening, the overall estimated resolution of the map reached 4.2 Å (FSC_0.143_). An additional map of the full tip, of overall lower resolution (FSC_0.143_ 3.88 Å) was also calculated in order to be able to fit the entire tip model but was not used for model building. Further image processing was necessary to obtain a map including the monomeric p143 protein; refined particles from the tip reconstruction were re-extracted (box size of 200x200 pixels^2^) and re-centred on the lower part of BHP_pb3_, on the side of which p143 is located. After re-classification/selection, symmetry relaxation and a new run of 3D refinement using suitable masking, a non-symmetrized map of the central part of the tip was obtained, where densities for the monomeric p143 are visible. ***Tip-FhuA***. After particle extraction (box size of 340x340 pixels^2^) and extensive 2D and 3D classification, a homogeneous dataset of 20,349 particles was obtained. As an initial model, an 8 Å resolution map (low-pass filtered at 15 Å) obtained from a previous cryo-EM data collection (G. Effantin, unpublished) was used and a reconstruction of the full non-symmetrized tip after interaction with FhuA-nanodisc (Full tip-FhuA) was calculated. After masking and sharpening, the overall estimated resolution of the map reached 4.3 Å (FSC_0.143_). Signal subtraction was then performed to enhance the resolution of specific parts of the structure. Based on the previously determined reconstruction, two soft masks were created, for the bent fibre only and for the C3 open tube. After re-extraction and 3D reconstruction, the overall quality of these two areas greatly improved, with overall estimated resolution of 4.3 Å and 3.60 Å respectively for the bent fibre and the open tube.

### Tip/Tip-FhuA common core

In order to improve the resolution of the tip common core (TTP_pb6_, p132, p140, DTP_pb9_), and because we observed that it was invariant whether or not the tails were incubated with FhuA-nanodisc, particles from all three datasets were merged. A soft mask was created using a 20 Å resolution model generated with Chimera tool molmap, using a previously built atomic model (see “Protein model building” below) containing two TTP_pb6_ trimers, a p140 trimer, a DTP_pb9_ hexamer and a p132 dodecamer and used to perform signal subtraction on the merged particles. Subtracted particles were then refined to obtain a better C3 reconstruction for the tip common core, whose resolution reached an overall estimated value of 3.4 Å, allowing to build slightly more accurate atomic models for TTP_pb6_, p140, DTP_pb9_ and p132 proteins. Efforts to specifically isolate and align the ill-defined RBP_pb5_/FhuA-RBP_pb5_ parts of the maps did not result in any improvement, probably due to the small size of the protein and/or the low number of particles.

For every reconstruction, a local resolution map was calculated using Relion built-in local resolution tool (Extended Data Fig. 1c, h).

### Protein model building

Atomic protein models were built into the different cryo-EM maps (Table S4) using the Coot software (46) by tracing the protein sequence into the densities and were then iteratively refined alternating Coot manual refinement and PHENIX (47) real space refine tool until convergence. p140, p132, BHP_pb3_ and TMP_pb2C_ models were built *ab initio*. For TTP_pb6_ and DTP_pb9_ models, existing X-ray models (5NGJ(14) / 4JMQ (17)) were used as a starting point and were refined into the EM maps. Molprobity (48) was used for model quality assessment. The densities corresponding to the BHP_pb3_ β-hairpin legs (residues ∼225-245) in the Tip-FhuA map are poorly resolved with regard to the rest of the protein. As a consequence, we only propose a likely model for BHP_pb3_ β-hairpins, which should be considered with caution.

For p143, we used an AlphaFold2 (31) predicted model as a starting point, which was fitted into the corresponding densities. Of note, Alphafold2’s level of confidence was not optimal throughout the whole sequence, which could explain the partial fit of the initial model. It was though coherent regarding the global shape and size. We then used a combination of Flex-EM (49) / Namdinator (50) (flexible fitting) and PHENIX (47) (real space refine), in an iterative way, to obtain a better model for this protein, with a convincing fit of most of its secondary structures.

### Proteomics based on High Performance Liquid Chromatography/Electrospray Ionization (HPLC/ESI)-orbitrap

T5 tail proteins were stacked in the top of a 4–12% NuPAGE gel (Invitrogen) and stained with R-250 Coomassie blue. Gel bands were manually excised and cut in pieces before being subjected to digestion using modified trypsin (Promega, sequencing grade) as previously described (51). Peptides were analysed by online nanoLC-MS/MS (UltiMate 3000 RSLCnano and QExactive Plus, Thermo Scientific) with 2 replicates per sample. Peptides were sampled on a 300 μm x 5 mm PepMap C18 precolumn and separated on a 75 μm × 250mm C18 column (PepMap, Dionex). The nanoLC method consisted of a 120-min gradient at a flow rate of 300 nL/min, ranging from 5 to 37% acetronitrile in 0.1% formic acid for 114 min, before reaching 72% acetronitrile in 0.1% formic acid for the last 6 min.

Spray voltage was set at 1.6 kV; heated capillary was adjusted to 270°C. Survey full-scan MS spectra (m/z = 400–1,600) were acquired with a resolution of 70,000 after accumulation of 10^6^ ions (max. fill time : 200 ms). The 10 most intense ions were fragmented by higher-energy collisional dissociation after accumulation of 10^5^ ions (max. fill time : 50 ms). LC-MS/MS datafiles were processed using MaxQuant, version 1.5.1.2 (52). Spectra were searched against the Uniprot database and frequently observed contaminants database. The I = L option was activated. Trypsin enzyme was selected and 2 missed cleavages were allowed. Peptide modifications: carbamidomethylation (C, fixed), acetyl (Protein N-ter, variable) and oxidation (M, variable). Minimum number of unique peptides: 1. Matching between runs option activated. The proteomics data were deposited onto the ProteomeXchange server^20^ with the dataset identifier PXD004215. Statistical analyses were performed using the Perseus toolbox (version 1.5.1.6) within MaxQuant. Proteins identified in the reverse and contaminant databases, or with less than 2 razor + unique peptides, or exhibiting less than 6 iBAQ values in one condition were discarded. After log2 transformation, iBAQ values for the 1952 remaining proteins were normalized by condition-wise centring, missing values were imputed for each injected sample as the 2.5-percentile value and statistical testing were conducted using welch t-testing.

### Liquid Chromatography/Electrospray-Ionization Mass Spectrometry (LC-ESI-TOF-MS)

To measure the accurate mass of the tail proteins, the T5 tails were diluted 2:3 in 0.1% TFA to obtained a final tail concentration of 0.20-0.22 μM and were analysed using Electrospray (ESI)-time of flight (TOF) mass spectrometer (6210 instrument, Agilent Technologies) coupled to a Liquid Chromatography (LC) system (1100 series, Agilent Technologies). The instrument was calibrated using tuning mix (ESI-L, Agilent Technologies). The following instrumental settings were used: gas (nitrogen) temperature 300 °C, drying gas (nitrogen) 7 L.min^−1^, nebulizer gas (nitrogen) 10 psig, Vcap 4 kV, fragmentor 250 V, skimmer 60 V, Vpp (octopole RF) 250 V. The HPLC mobile phases were prepared with HPLC grade solvents. Mobile phase A composition was: 5% ACN, 0.03% TFA. Mobile phase B composition was: 95% ACN, 0.03% TFA.

8 μl of each sample (1.6-1.8 pmol) were first desalted on-line for 3 min with 100% of mobile phase A (flow rate of 50 μl / min), using a C8 reverse phase micro-column (Zorbax 300S**B-**C8, 5μm, 5 × 0.3 mm, Agilent Technologies). The sample was then eluted with 70% of mobile phase B (flow rate of 50 μl/ min) and MS spectra were acquired in the positive ion mode in the 300-3000 m/z range. Data were processed with MassHunter software (v. B.02.00, Agilent Technologies) and GPMAW software (v. 7.00b2, Lighthouse Data, Denmark).

## Supplementary information for

### Other Supplementary Materials for this manuscript include the following

**Supplementary Movie S1 and S2: Morphs of the BHP**_**pb3**_ **trimer between the conformation before and after interaction with the receptor**. Side (S1) and top (S2) views. Monomers are coloured yellow, orange and salmon, with the HDIV-FNIII linker and the two FNIII domains in darker shades of the same colour.

**Supplementary Movie S3 and S4**: **Morphs of the BHP**_**pb3**_ **trimer without its FNIIIs, between the conformation before and after interaction with the receptor**. Side (S3) and top (S4) views. Monomers are coloured yellow, orange and salmon, with the HDI and DHIV domains in lighter colours, the HDIV-FNIII linker in dark orange and the plugs in brown.

**Figure S1:**
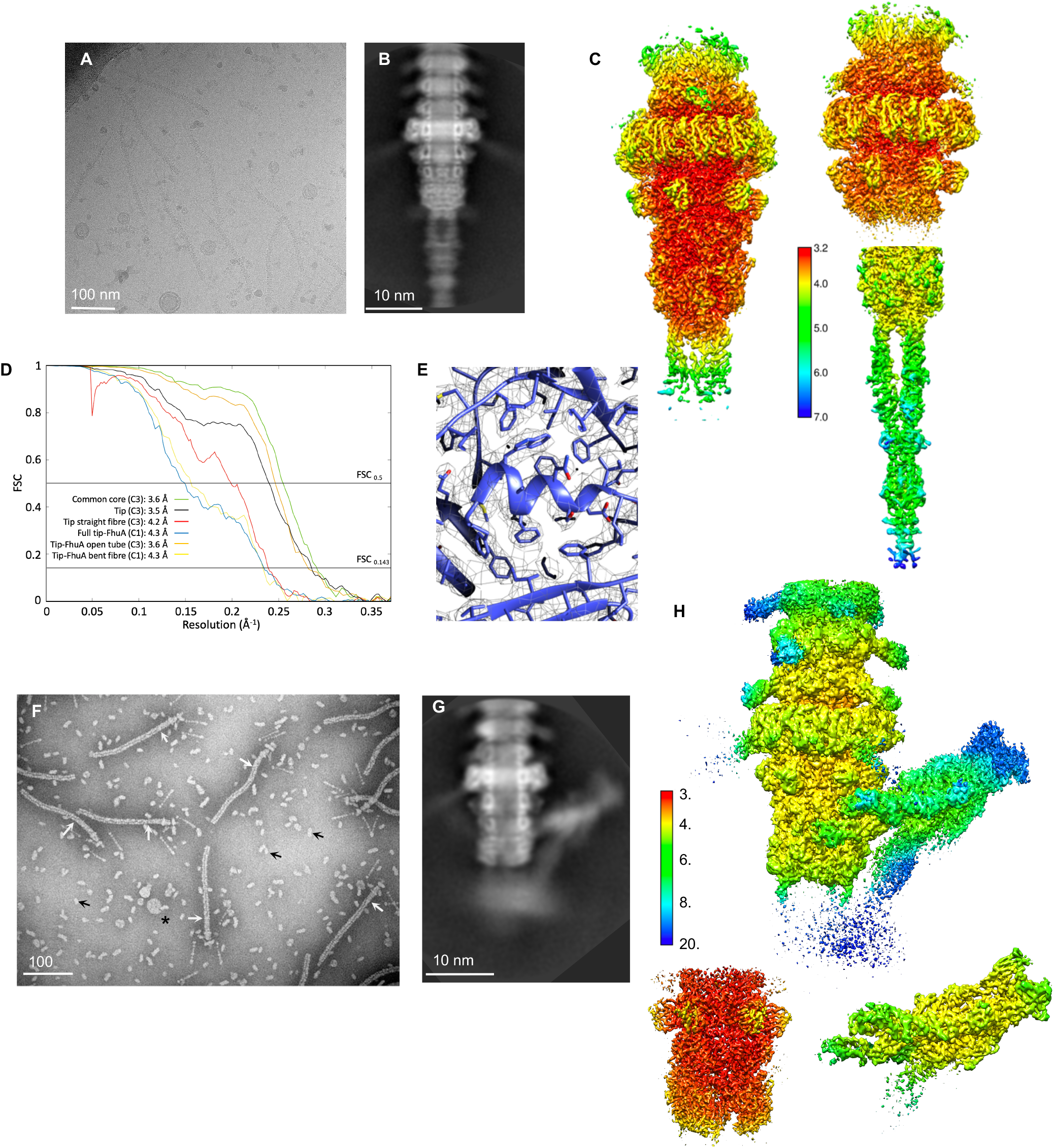
**A**. Cryo-EM image of T5 tails. **B**. A 2D class average of T5 tail tip. **C**. Local resolution maps, as determined by Relion of the tip (left), the Tip/Tip-FhuA common core (top right, from TTP_pb6_ to the BHP_pb3_ top part, map obtained gathering particles from both Tip and Tip-FhuA datasets, top right) and of the tip straight central fibre (bottom right). All maps were calculated with a C3 symmetry and the key (in Å) is the same for the three maps. Resolution is the highest at the centre (3.2 Å) and falls off radially to ∼7 Å at the tip of TTP_pb6_ Ig-like domain. Because of the flexibility of the central fibre and of the tube, resolution also drops rapidly along the tube, above the first TTP_pb6_ ring and downwards along the central fibre. **D**. Fourier shell correlation plot for the six maps presented in C. and H. FSC_0.5_ and FSC_0.143_ cutoffs are indicated, as well as the estimated resolution (FSC_0.143_) for each map. **E** Close up view of an α-helix of p140 monomer model, built using Tip/Tip-FhuA common core map. **F**. Large field, negative stain EM image of T5 tails incubated with FhuA-nanodiscs. The background is filled with FhuA-nanodiscs, mainly seen lying on the side (black arrows) but also from the top (*). T5 tails are partially emptied from TMP_pb2_. White arrows point to the empty/filled limit in the tails. **G**. A 2D class average of Tip-FhuA. (**H)** Local resolution maps, as determined by Relion, of the C1 full Tip-FhuA (left), C1 bent fibre (BHP_pb3_ FNIII, pb4, top right) and C3 open tube (DTP_pb9_, BHP_pb3_ ∼700 N-terminal residues, TMP_pb2*_ 43 C-terminal residues, bottom right). The key (in Å) is the same for the three maps.

**Figure S2:**
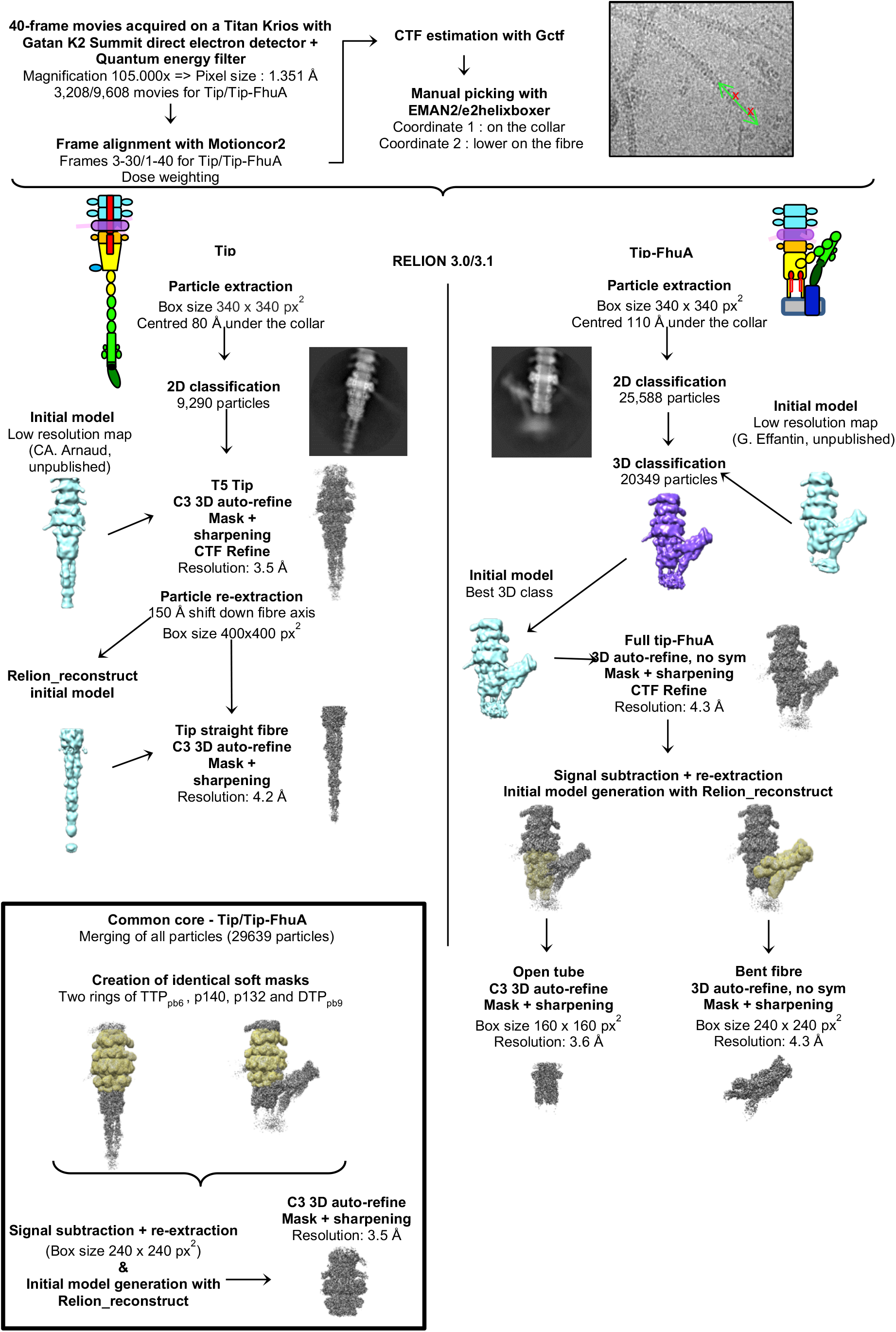
Flowchart of the EM processing pipeline for T5 Tip and Tip-FhuA. Common steps for the Tip/Tip-FhuA datasets are framed in blue. Initial models are in blue, 3D classes in purple, soft masks in yellow and 3D refine in grey. Top right: vectorial picking of the tip particles on a cryo-EM image of a T5 tail.

**Figure S3:**
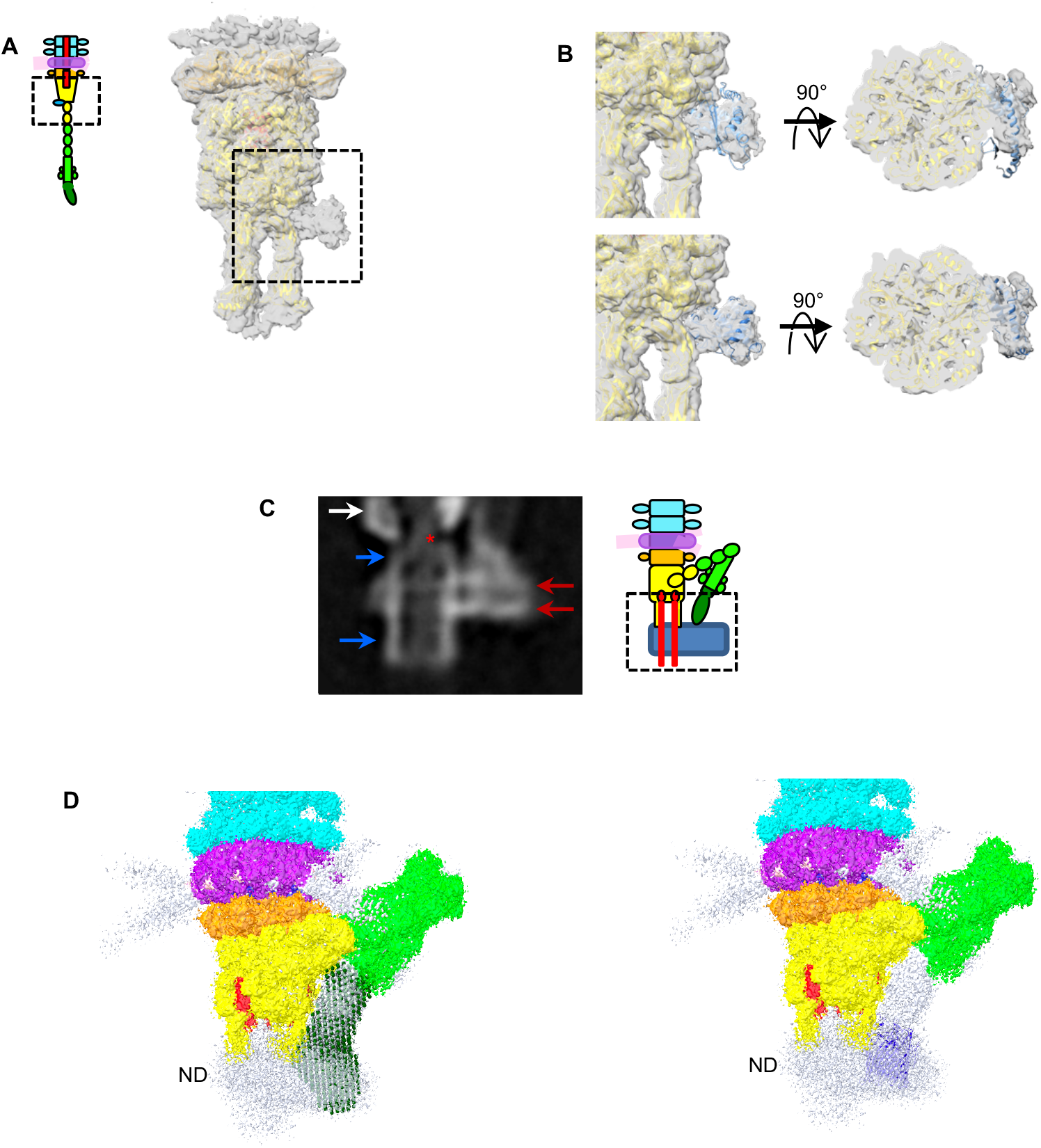
**A**. Isosurface view of an unsymmetrised cryo-EM map of the tip, with models for the BHP_pb3_ trimer (yellow) and the DTP_pb9_ hexamer (orange) fitted in it. Additional densities at the base of BHP_pb3_, corresponding to a monomeric protein, are clearly visible (dotted box). **B**. Enlargement of the dotted box in a, with a fit of the Alphafold2 prediction structure for p143 before (top, light blue model) and after (bottom, dark blue model) flexible fitting. Left: side view, right: top view from a slice. **C**. 2D projection of a low resolution EM reconstruction from Tip-FhuA showing the bottom of BHP_pb3_ (white arrow), the nanodisc bilayer (red arrows) and what appears to be cylindrical structure connected to BHP_pb3_ and spanning the nanodisc (blue arrows), probably the channel or part of it. Densities corresponding to TMP_pb2*_ can also been seen inside the tube/channel (red asterisk). **D**. Left: Fit of a SANS envelop of the FhuA-pb5 complex (dark green beads) into the densities prolonging pb4 spike and merging into the nanodisc. Right: Fit of FhuA (PDB 2GRX, blue ribbon) in the same map, in the nanodisc below the RBP_pb5_ density. The map is that of Tip-FhuA C1 filtered and masked reconstruction, coloured as in figure 1 and 40% transparent. Unattributed densities are white.

**Figure S4:**
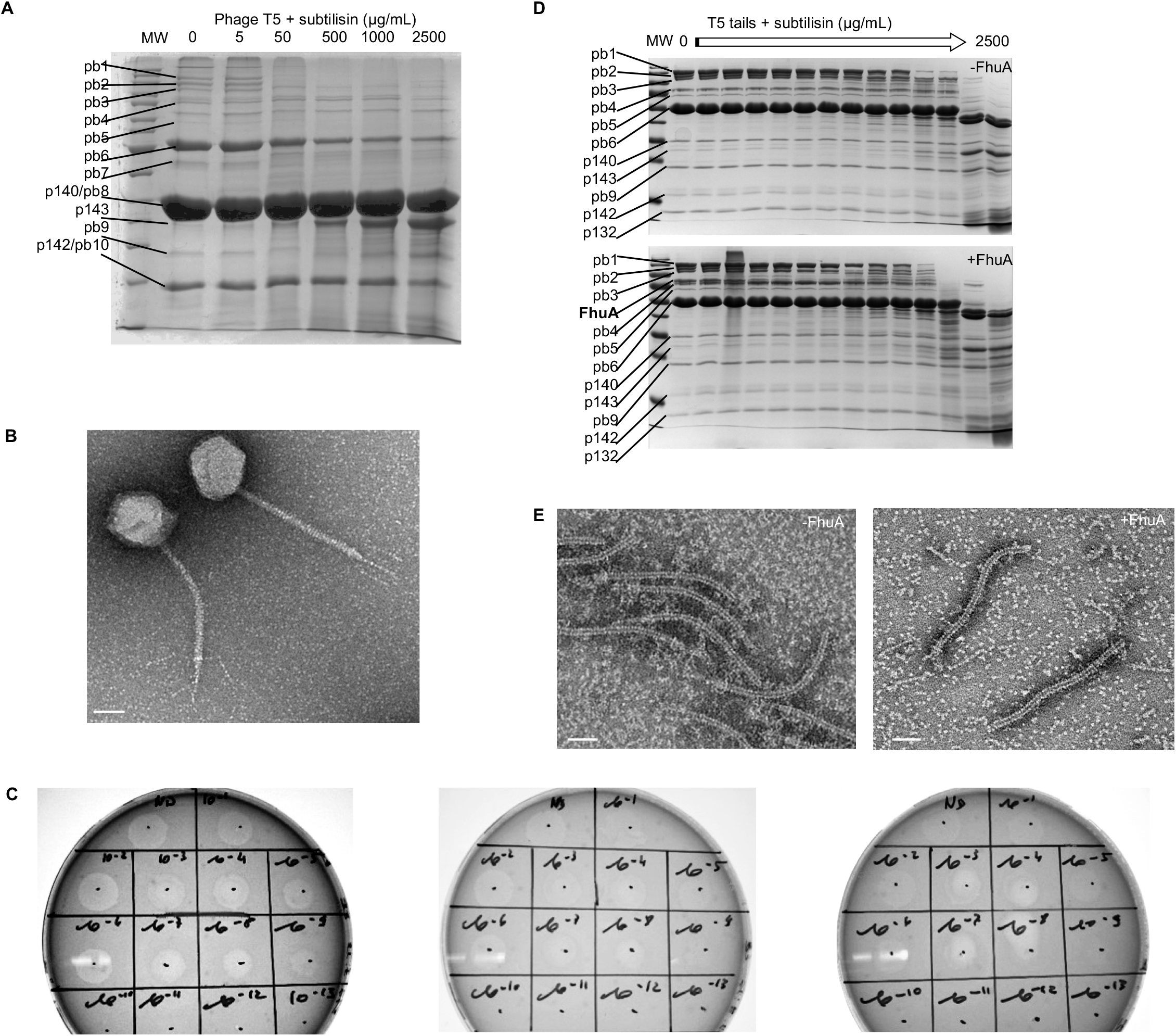
Limited proteolysis of phage T5 (A, B, C) and T5 tails (D, E). **A**. Phage T5 was incubated with subtilisin (0, 5, 50, 500, 1000, 2500 μg/mL) for 1h at room temperature. The reaction was stopped by the addition of 10 mM PMSF, and phage ghosts (*11*) were prepared for migration on 12% SDS-PAGE. Molecular weight markers: 200, 150, 120, 100, 85, 70, 60, 50, 40, 30, 25, 20, 15, 10 kDa. **B**. Negative stain EM image of phage T5 after 1h incubation with 2.5 mg/mL subtilisin. **C**. Titration of unproteolysed T5 (left), T5 incubated 1h with 50 μg/mL (middle) and 2.5 mg/mL (right) subtilisin on a lawn of *E. coli* strain F in soft Agar. **D**. Upper panel: Purified T5 tails were incubated with subtilisin (0, 0.1, 0.3, 0.6, 1, 2, 3, 5, 7.5, 10, 20, 30, 1000, 2500 μg/mL). Lower panel: T5 tails were pre-incubated with LDAO-solubilised FhuA for 1h at room temperature (final LDAO concentration 0.05%), before being incubated with subtilisin. Molecular weight markers: 180, 130, 100, 70, 55, 40, 35, 25, 15, 10 kDa. **E**. Negative stain EM images of T5 tails (right) and T5 tails incubated with FhuA (left) incubated with 2.5 mg/mL subtilisin. Subtilisin is clearly visible in the background. Bar: 50 nm.

**Figure S5:**
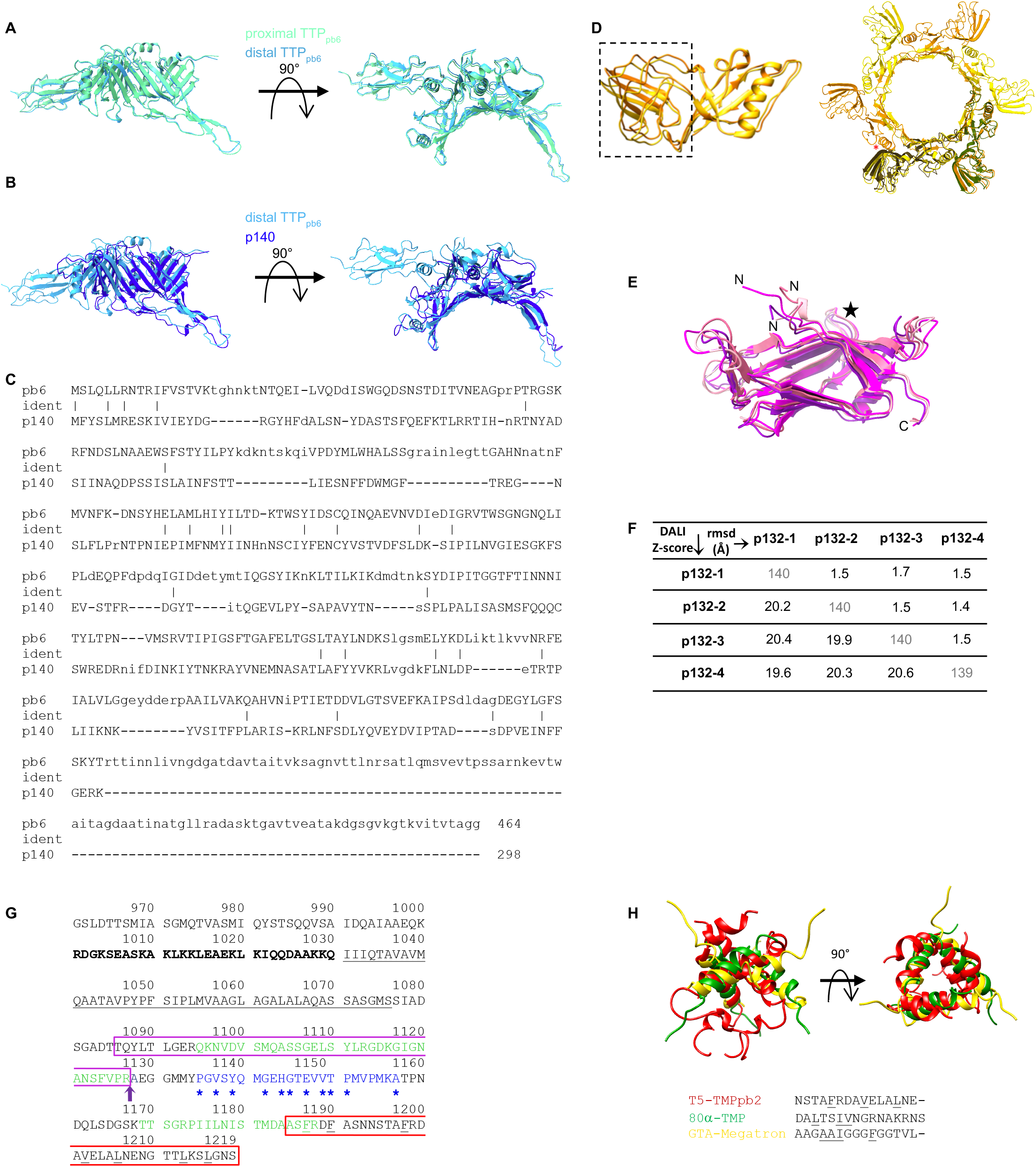
Comparison between upper and lower TTP_pb6_ (**A**) and between lower TTP_pb6_ and p140 (DALI Z-score of 22.6, with an rmsd over 282 residues of 2.6 Å) (**B**). Side (left) and top (right) views. **C**. Sequence alignment between TTP_pb6_ and p140. Residues from aligned structures are uppercase. Identical residues are highlighted with a vertical bar. **D**. Left: Superimposition of the two DTP_pb9_ subunits unrelated by symmetry, aligned on the TTD (rmsd 1.1 Å over 204 residues TTD 0.8 Å over 112 residues), the OB domain is framed by a rectangle. Right: DTP_pb9_ hexameric ring, in which two subunits of the asymmetric unit are superimposed, on the TTD, with the crystal structure of DTP_pb9_ (in green). The OB domain of monomer 2 is making contact with the TTD of monomer 1 (*). **E**. DALI superposition of the four p132 of the asymmetric unit. The N- and C-termini are indicated and the star marks loop 52-60. **F**. Table summarising the pairwise superpositions of the four p132 subunits of the asymmetric unit. In the diagonal in grey is the number of residues built for each subunit, above the diagonal the rmsd in Å and below the DALI Z-score. DALI search links the p132 fold to the N-terminal domain of the Baseplate Protein Upper (BppU, ORF48) of phage TP901-1 with a Z-score of 5.9 and a rmsd of 4.0 Å over 99 residues and 9% identity. **G**. Sequence analysis of the TMP_pb2_ 259 C-terminal residues. Bold: predicted coiled-coil region from COILS; underlined: predicted transmembrane domain from PSI-PRED and TMHMM; green: peptides identified in proteomics; blue: Zinc carboxypeptidase motif, with conserved residues from Prosite analysis indicated with a star; purple arrow: cleavage site, separating TMP_pb2*_ and TMP_pb2C_. The red box corresponds to TMP_pb2C_ 35 modelled residues from the Tip map, those pointing towards the centre of the coil are underlined. The magenta box correspond to TMP_pb2*_ C-terminus 42 modelled residues in Tip-FhuA map. **H**. Overlay of T5 TMP_pb2C_ (red), TMP_80a_ (green) and the N-terminal helix of GTA Megatron (yellow) when superimposing the BHP trimers of each baseplate. Sequence alignment of the aligned helices, with the hydrophobic residues pointing towards the centre of the coil underlined.

**Figure S6:**
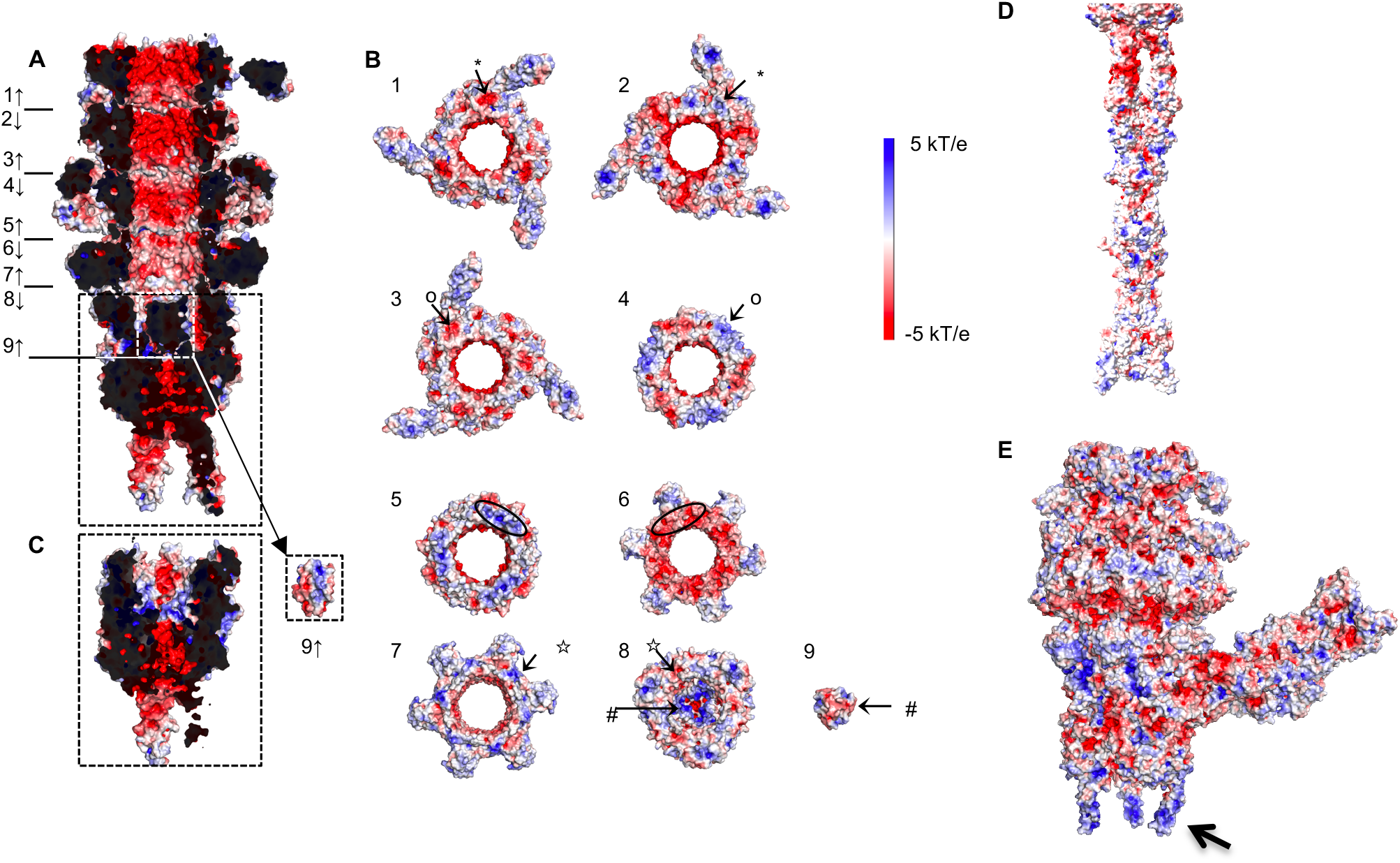
Electrostatic charge distribution of **A**. the interior of T5 tail tube, **B**. the complementary interfaces of the different rings, **C**. the closed BHP_pb3_ in which TMP_pb2c_ has been removed *in silico* and is shown on the right side, **D**. the tail tip before and **e** after interaction with FhuA. The arrow points to one of the highly charged three β-hairpins “legs”. The position of the views in **B** is indicated by a number in **A**, and complementary charge patches are noted by symbols. Electrostatic charge distribution was calculated from the APBS plugin of either PyMol or ChimeraX.

**Figure S7:**
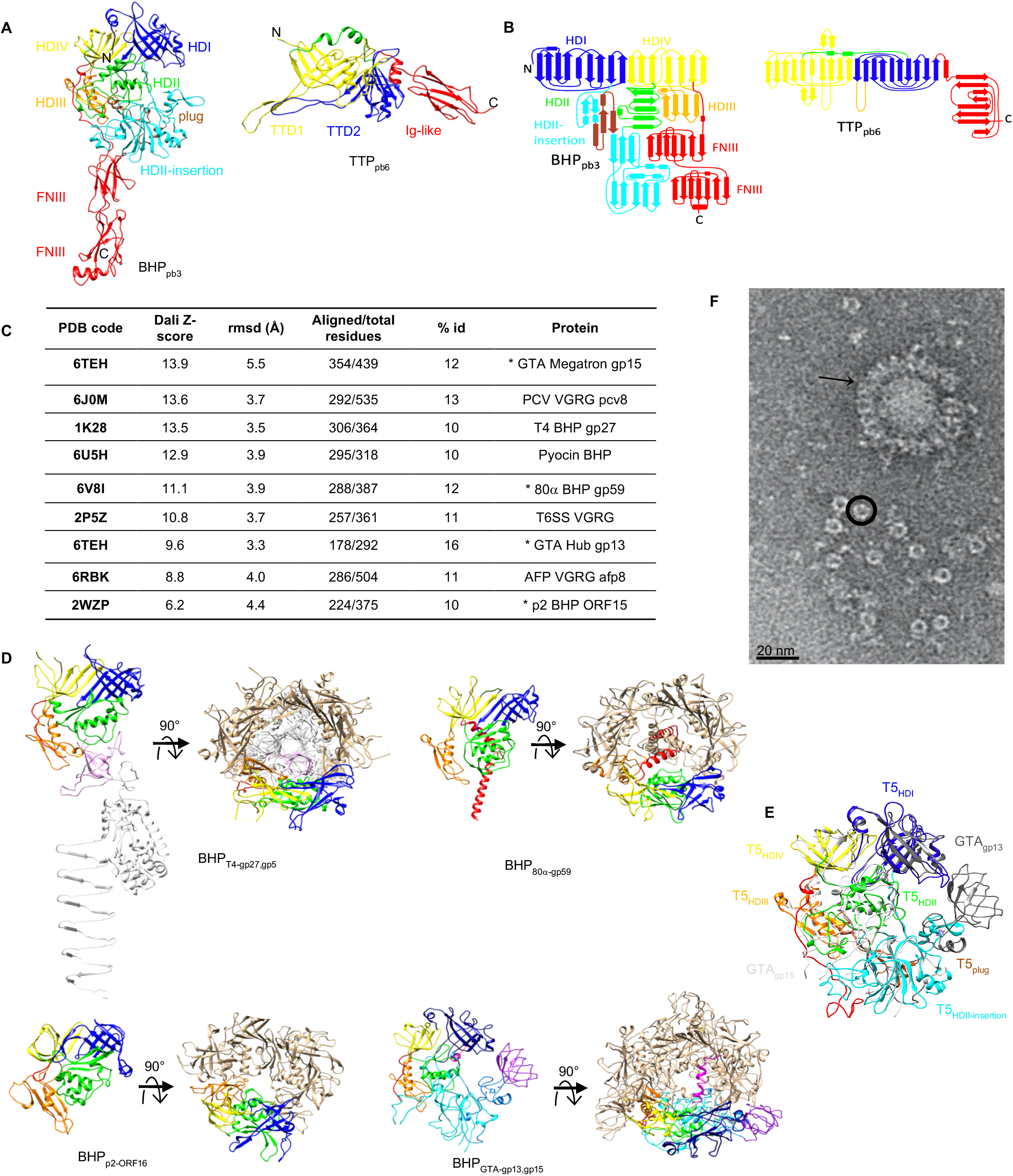
**A**. Ribbon representation in the same side view orientation of TTP_pb6_ (left) and BHP_pb3_ (right) and **B**. topology diagram of the two proteins. BHP_pb3_ HDI-VI are coloured blue, green, orange and yellow, respectively, the HDII insertion in cyan and the two FNIII in red. The same colour code was used for structurally homologous domain in TTP_pb6_. The link between the two Tail Tube domains (TTD) in BHP_pb3_ and TTP_pb6_ is topologically different in the two proteins: it connects the C-terminus of TTD1 domain to the N-terminus of TTD2 in TTP_pb6_, and the N-terminus TTD1 domain to the C-terminus of TTD2 in BHP_pb3_. N- and C-terminus are indicated. **C**. DALI search using BHP_pb3_ without its FNIII (716 residues) as a bait. Asterix point to *Siphoviridae* BHPs. **D**. Side views of monomeric BHP proteins of T4-gp27, 80α-gp59, p2-ORF16 and GTA-gp13-15 with the same colour code as in A, and top views of the trimeric complex, with one monomer coloured. In T4, gp5 was added, with the OB domain that closes the tube coloured in pink and the needle in white. Note that GTA BHP is composed of two proteins, the Hub (gp13) and the Megatron (gp15). The Hub protein comprises HDI (darker blue), part of the insertion domain (darker cyan) in which is inserted an OB domain (purple), while the Megatron comprises HDII (green), part of the insertion domain (cyan), HDIII (orange), HDIV (yellow) and an N-terminal “iris helix” (magenta). **E**. Side view superimposition of BHP_pb3_ (coloured) with GTA-Megatron (light grey) and GTA-Hub (dark grey) in ribbon representation. **F**. Negative stain EM images of purified BHP_pb3_ showing isolated monomers coexisting with either free trimers seen in top views (circled), or that aggregate around impurities, seen in side views (arrow). The dimensions of the trimer (hight: 45 nm, diameter: 9 nm) correspond to those of the BHP_pb3_ trophy cup. In the top views, six subdomains can be distinguished (see circled particle).

**Figure S8:**
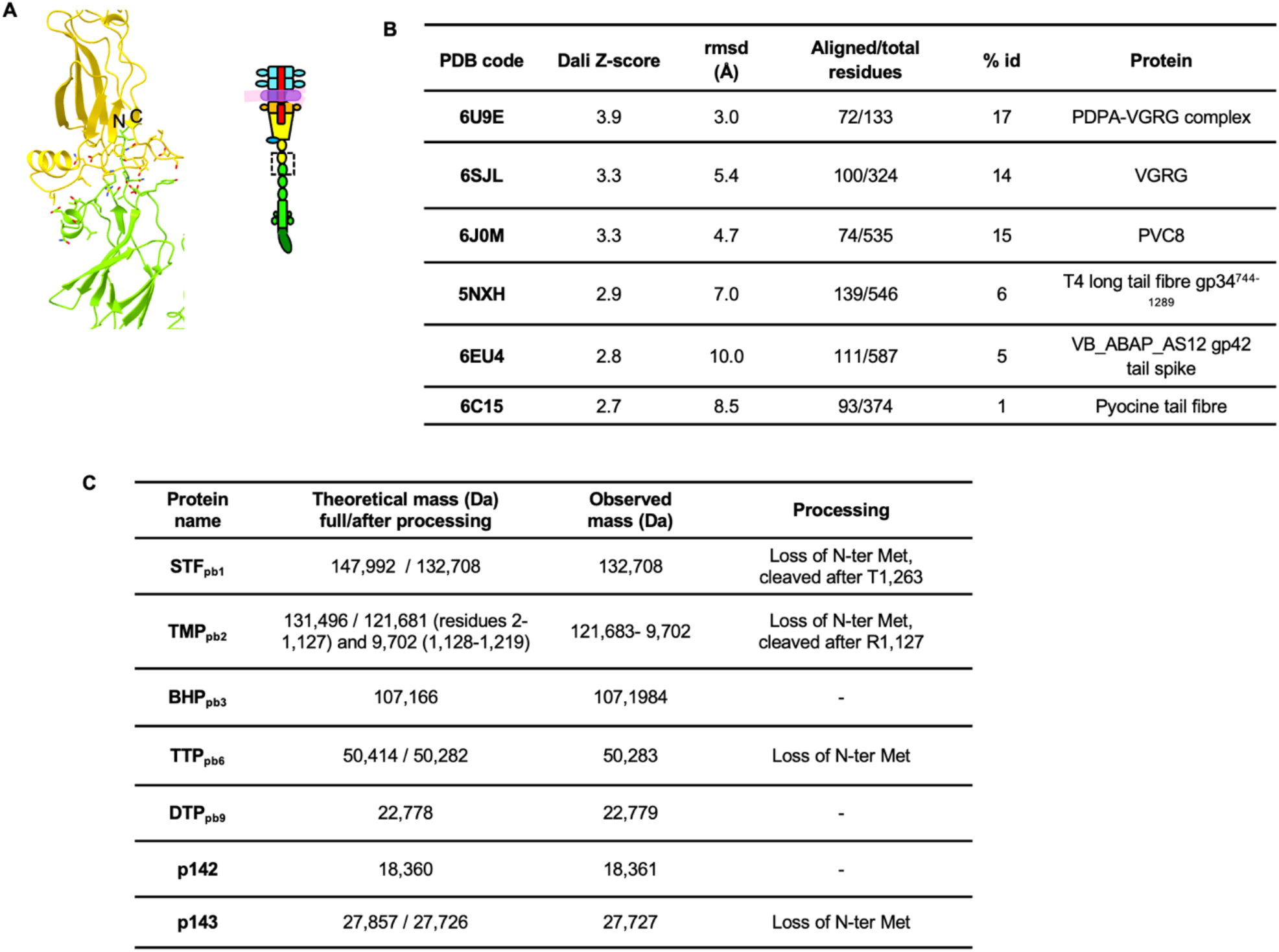
**A**. interface between BHP_pb3_ last FNIII (yellow) and pb4 first FNIII (green) in ribbon representation. BHP_pb3_ C-terminus (C) and pb4 N-terminus (N) are highlighted. Residues at the interface between the two proteins are shown in sticks. **B**. Table of relevant DALI hits for pb4 spike. Pb4 domain 2 is linked to phage spike decoration domains (PDB codes 7CHU-A, 6TGF-D, 6E1R-A, 5M9F-A, 6NW9-C, 6EU4-A, 5W6H-A with DALI Z-score of 3.9 to 2.3, rmsd from 2.4 to 3.0 Å over ∼45 residues and identity ranging from 4 to 14%). **C**. LC-ESI-TOF-MS of purified tails. The experimental intact mass of seven T5 tail proteins were determined. Even though proteomics detected their presence (Table S3), masses of pb4, RBP_pb5_, p140 and p132 as intact proteins could not be measured. This may be explained by a difficulty in their ionisation.

**Table S1:**
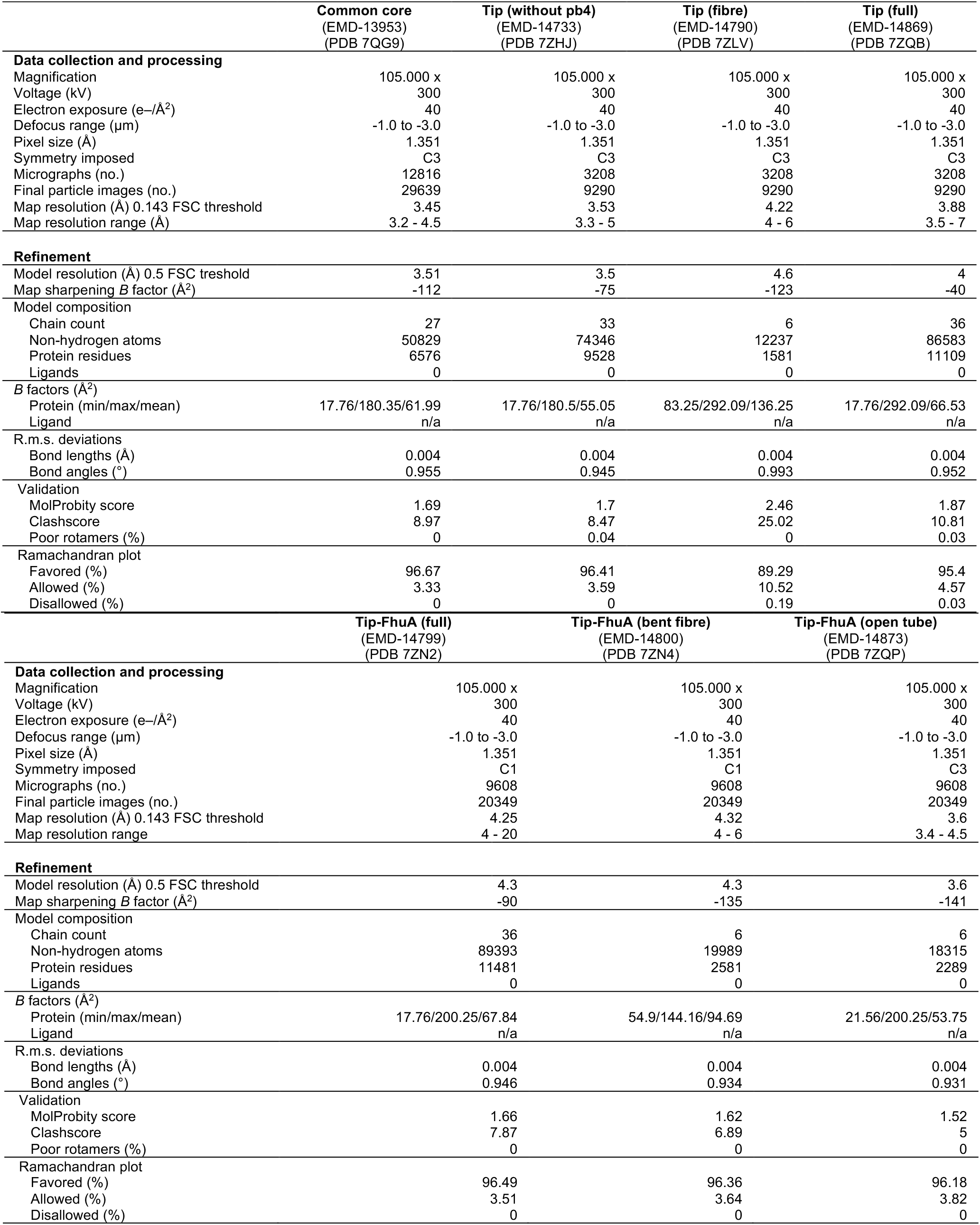
Cryo-EM data collection, refinement and validation statistics.

**Table S2:** Domain definition of T5 tip proteins. STF_pb1_, RBP_pb5_ and p143, in italics, are visible but partially or not resolved in our EM maps. The colour code is that of Figure 1.

| **T5 protein name /**<br>**function** | **Number of**<br>**residues** | **Domains** | **Sequence** |
| --- | --- | --- | --- |
| TPP<sub>pb6</sub><br>Tail Tube Protein | 464 | TTD1<br>TTD2<br>Ig-Like | 1-203<br>204-355<br>356-464 |
| p140<br>Baseplate Tube Protein | 298 | TTD1<br>TTD2 | 1-167<br>168-298 |
| p132<br>Collar protein | 140 | Ig-like | 1-140 |
| STF<sub>pb1</sub><br>Side Tail Fibre | 1,263 |  |  |
| DTP<sub>pb9</sub><br>Distal Tail Protein | 204 | TTD<br>OB | 1-83; 172-204<br>84-171 |
| BHP<sub>pb3</sub><br>Baseplate Hub Protein | 949 | HDI<br>HDII-Insertion<br>plug/β-hairpin<br>DHII<br>HDIII<br>HDIV<br>HDIV-FNIII linker<br>FNIII-1<br>FNIII-2 | 1-159<br>160-437<br>210-262<br>438-570<br>586-658<br>571-585; 659-709<br>711-742<br>743-835<br>836-949 |
| TMP<sub>pb2</sub><br>Tape Measure Protein | 1,219* |  | 2-1127, 1128-1219 |
| pb4<br>Central Fibre Protein | 688 | FNIII-1<br>FNIII-2<br>FNIII-3<br>FNIII-spike linker<br>Spike<br>Small domains | 1-105<br>106-208<br>209-316<br>317-332<br>333-465; 627-688<br>466-626 |
| RBP<sub>pb5</sub><br>Receptor Binding Protein | 640 |  |  |
| p143<br>Tail Completion Protein | 262 |  |  |

**Table S3:** Proteomics of T5 tail proteins.

| Protein | Accession number | Gene name | Protein set score | Theoretical mass (Da) of full protein | #observable peptides | Coverage % |
| --- | --- | --- | --- | --- | --- | --- |
| **STF<sub>pb1</sub>** | **FIBL1_BPT5** | **ltf** | **8529.02** | **147992** | **81** | **73.14** |
| **TMP<sub>pb2</sub>** | **TMP_BPT5** | **D18-19** | **7609.99** | **131496** | **75** | **62.84** |
| **BHP<sub>pb3</sub>** | **BPPB3_BPT5** | **D16** | **4554.46** | **107166** | **53** | **75.66** |
| **pb4** | **FIBC_BPT5** | **D17** | **3347.97** | **74788** | **31** | **68.02** |
| **RBP<sub>pb5</sub>** | **RBP5_BPT5** | **oad** | **2063.73** | **68726** | **25** | **54.38** |
| **TTP<sub>pb6</sub>** | **TUBE_BPT5** | **N4** | **5054.41** | **50414** | **22** | **95.69** |
| **p143** | **COMPL_BPT5** | **ORF136** | **970.79** | **27857** | **18** | **63.53** |
| **p140** | **TAIL1_BPT5** | **ORF133** | **1221.08** | **34334** | **12** | **62.75** |
| **DTP<sub>pb9</sub>** | **DIT_BPT5** | **D16** | **1315.54** | **22778** | **11** | **85.29** |
| **p132** | **FIBL2_BPT5** | **ORF125** | **375.13** | **15067** | **5** | **57.86** |
| **p142** | **TTTP_BPT5** | **ORF135** | **650.65** | **18360** | **5** | **40.37** |
| MCP<sub>pb8</sub> | CAPSD_BPT5 | D20 | 537.11 | 50885 | 31 | 24.45 |
| pb10 | DECO_BPT5 | N5 | 84 | 17247 | 11 | 9.15 |
| putative metallopeptidase/<br>ribonuclease | Q5DML2_BPT5 | ORF082 | 66.64 | 25001 | 12 | 10.36 |
| Uncharacterized protein | Q66M03_BPT5 | T5p046 | 256.58 | 27425 | 16 | 26.29 |
| Glucose-1-phosphatase | AGP_ECOLI | agp | 114.46 | 45683 | 24 | 6.05 |
| Elongation factor Tu | EFTU1_ECOLI | tufA | 86.7 | 43284 | 26 | 4.31 |
| Uncharacterized protein | YAGL_ECOLI | yagL | 29.97 | 27274 | 16 | 3.88 |
| Glyceraldehyde-3-phosphate<br>dehydrogenase A | G3P1_ECOLI | gapA | 27.76 | 35532 | 24 | 3.32 |
| Uncharacterized protein | YDHW_ECOLI | ydhW | 26.29 | 24421 | 13 | 4.19 |

Bold proteins are tail proteins (11). MCP<sub>pb8</sub> and pb10 are capsid proteins, identified because of a small proportion of revertant ( $10^{-7}$ ) in the T5D20am30d mutant, leading to the production of full T5 particles.
Q66M03\_BPT5: uncharacterised T5 protein, gene located in the early gene region. It is surrounded by genes coding for lysins, holins and endolysins.
Q5DML2\_BPT5: T5 protein, annotated as a putative metallopeptidase/carboxypeptidase. HHPred aligns it with nucleases or a subunit of DNA polymerase III. It is located in the early gene portion of the genome, surrounded by DNA and RNA interacting proteins. Phagocyte (53) aligns it with similar proteins in T5-like phages, but also other siphon- and myo-phages.

**Table S4:**
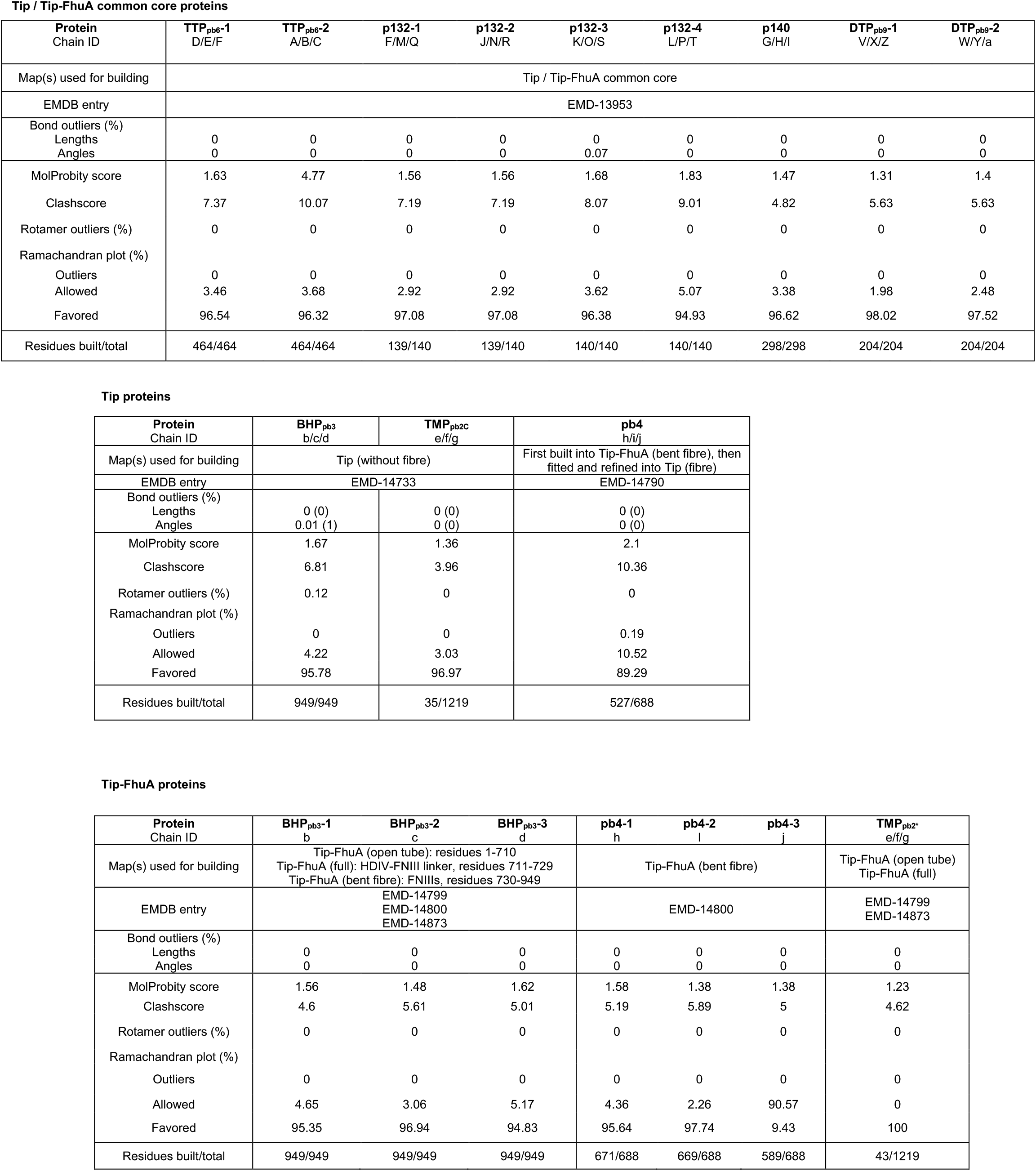
Validation statistics and model building for T5 tip individual proteins.

## References

1. C. A. Suttle, Marine viruses--major players in the global ecosystem. Nat. Rev. Microbiol. 5, 801–812 (2007).

2. S. Uyttebroek, B. Chen, J. Onsea, F. Ruythooren, Y. Debaveye, D. Devolder, I. Spriet, M. Depypere, J. Wagemans, R. Lavigne, J.-P. Pirnay, M. Merabishvili, P. De Munter, W. E. Peetermans, L. Dupont, L. Van Gerven, W.-J. Metsemakers, Safety and efficacy of phage therapy in difficult-to-treat infections: a systematic review. Lancet Infect Dis, S1473-3099(21)00612–5 (2022).

3. D. Veesler, C. Cambillau, A Common Evolutionary Origin for Tailed-Bacteriophage Functional Modules and Bacterial Machineries. Microbiol Mol Biol Rev. 75, 423–433 (2011).

4. A. R. Davidson, L. Cardarelli, L. G. Pell, D. R. Radford, K. L. Maxwell, Long noncontractile tail machines of bacteriophages. Adv. Exp. Med. Biol. 726, 115–142 (2012).

5. M. Brackmann, S. Nazarov, J. Wang, M. Basler, Using Force to Punch Holes: Mechanics of Contractile Nanomachines. Trends Cell Biol. 27, 623–632 (2017).

6. R. Linares, C.-A. Arnaud, S. Degroux, G. Schoehn, C. Breyton, Structure, function and assembly of the long, flexible tail of siphophages. Curr Opin Virol. 45, 34–42 (2020).

7. A. Desfosses, H. Venugopal, T. Joshi, J. Felix, M. Jessop, H. Jeong, J. Hyun, J. B. Heymann, M. R. H. Hurst, I. Gutsche, A. K. Mitra, Atomic structures of an entire contractile injection system in both the extended and contracted states. Nat Microbiol. 4, 1885–1894 (2019).

8. F. Jiang, N. Li, X. Wang, J. Cheng, Y. Huang, Y. Yang, J. Yang, B. Cai, Y.-P. Wang, Q. Jin, N. Gao, Cryo-EM Structure and Assembly of an Extracellular Contractile Injection System. Cell. 177, 370-383.e15 (2019).

9. P. Ge, D. Scholl, N. S. Prokhorov, J. Avaylon, M. M. Shneider, C. Browning, S. A. Buth, M. Plattner, U. Chakraborty, K. Ding, P. G. Leiman, J. F. Miller, Z. H. Zhou, Action of a minimal contractile bactericidal nanomachine. Nature. 580, 658–662 (2020).

10. H.-W. Ackermann, Phage classification and characterization. Methods Mol. Biol. 501, 127–140 (2009).

11. Y. Zivanovic, F. Confalonieri, L. Ponchon, R. Lurz, M. Chami, A. Flayhan, M. Renouard, A. Huet, P. Decottignies, A. R. Davidson, C. Breyton, P. Boulanger, Insights into bacteriophage T5 structure from analysis of its morphogenesis genes and protein components. J. Virol. 88, 1162–1174 (2014).

12. M. Demerec, U. Fano, Bacteriophage-Resistant Mutants in Escherichia Coli. Genetics. 30, 119–136 (1945).

13. A. Huet, R. L. Duda, P. Boulanger, J. F. Conway, Capsid expansion of bacteriophage T5 revealed by high resolution cryoelectron microscopy. Proc. Natl. Acad. Sci. U.S.A. 116, 21037–21046 (2019).

14. C.-A. Arnaud, G. Effantin, C. Vivès, S. Engilberge, M. Bacia, P. Boulanger, E. Girard, G. Schoehn, C. Breyton, Bacteriophage T5 tail tube structure suggests a trigger mechanism for Siphoviridae DNA ejection. Nat Commun. 8, 1953 (2017).

15. P. Boulanger, P. Jacquot, L. Plançon, M. Chami, A. Engel, C. Parquet, C. Herbeuval, L. Letellier, Phage T5 straight tail fiber is a multifunctional protein acting as a tape measure and carrying fusogenic and muralytic activities. J Biol Chem. 286, 13556–13564 (2008).

16. C. Garcia-Doval, J. R. Castón, D. Luque, M. Granell, J. M. Otero, A. L. Llamas-Saiz, M. Renouard, P. Boulanger, M. J. van Raaij, Structure of the Receptor-Binding Carboxy-Terminal Domain of the Bacteriophage T5 L-Shaped Tail Fibre with and without Its Intra-Molecular Chaperone. Viruses. 7, 6424–6440 (2015).

17. A. Flayhan, F. M. D. Vellieux, R. Lurz, O. Maury, C. Contreras-Martel, E. Girard, P. Boulanger, C. Breyton, Crystal Structure of pb9, the Distal Tail Protein of Bacteriophage T5: a Conserved Structural Motif among All Siphophages. J. Virol. 88, 820–828 (2014).

18. A. Flayhan, F. Wien, M. Paternostre, P. Boulanger, C. Breyton, New insights into pb5, the receptor binding protein of bacteriophage T5, and its interaction with its Escherichia coli receptor FhuA. Biochimie. 94, 1982–1989 (2012).

19. C. Breyton, A. Flayhan, F. Gabel, M. Lethier, G. Durand, P. Boulanger, M. Chami, C. Ebel, Assessing the Conformational Changes of pb5, the Receptor-binding Protein of Phage T5, upon Binding to Its Escherichia coli Receptor FhuA. J. Biol. Chem. 288, 30763–30772 (2013).

20. V. Braun, FhuA (TonA), the career of a protein. J. Bacteriol. 191, 3431–3436 (2009).

21. M. Bonhivers, A. Ghazi, P. Boulanger, L. Letellier, FhuA, a transporter of the Escherichia coli outer membrane, is converted into a channel upon binding of bacteriophage T5. EMBO J. 15, 1850–1856 (1996).

22. N. Chiaruttini, M. de Frutos, E. Augarde, P. Boulanger, L. Letellier, V. Viasnoff, Is the in vitro ejection of bacteriophage DNA quasistatic? A bulk to single virus study. Biophys. J. 99, 447–455 (2010).

23. H. Fraga, C.-A. Arnaud, D. F. Gauto, M. Audin, V. Kurauskas, P. Macek, C. Krichel, J.-Y. Guan, J. Boisbouvier, R. Sprangers, C. Breyton, P. Schanda, Solid-State NMR H-N-(C)-H and H-N-C-C 3D/4D Correlation Experiments for Resonance Assignment of Large Proteins. Chemphyschem. 18, 2697–2703 (2017).

24. L. Holm, DALI and the persistence of protein shape. Protein Sci. 29, 128–140 (2020).

25. N. M. I. Taylor, N. S. Prokhorov, R. C. Guerrero-Ferreira, M. M. Shneider, C. Browning, K. N. Goldie, H. Stahlberg, P. G. Leiman, Structure of the T4 baseplate and its function in triggering sheath contraction. Nature. 533, 346–352 (2016).

26. J. L. Kizziah, K. A. Manning, A. D. Dearborn, T. Dokland, Structure of the host cell recognition and penetration machinery of a Staphylococcus aureus bacteriophage. PLoS Pathog. 16, e1008314 (2020).

27. D. Veesler, S. Spinelli, J. Mahony, J. Lichière, S. Blangy, G. Bricogne, P. Legrand, M. Ortiz-Lombardia, V. Campanacci, D. van Sinderen, C. Cambillau, Structure of the phage TP901-1 1.8 MDa baseplate suggests an alternative host adhesion mechanism. Proc Natl Acad Sci U S A. 109, 8954–8958 (2012).

28. G. Sciara, C. Bebeacua, P. Bron, D. Tremblay, M. Ortiz-Lombardia, J. Lichière, M. van Heel, V. Campanacci, S. Moineau, C. Cambillau, Structure of lactococcal phage p2 baseplate and its mechanism of activation. Proc Natl Acad Sci U S A. 107, 6852–6857 (2010).

29. E. Krissinel, K. Henrick, Inference of macromolecular assemblies from crystalline state. J. Mol. Biol. 372, 774–797 (2007).

30. M. Noirclerc-Savoye, A. Flayhan, C. Pereira, B. Gallet, P. Gans, C. Ebel, Cécile Breyton, Tail proteins of phage T5: investigation of the effect of the His6-tag position, from expression to crystallisation. Protein Expr. Purif. 109, 70–78 (2015).

31. J. Jumper, R. Evans, A. Pritzel, T. Green, M. Figurnov, O. Ronneberger, K. Tunyasuvunakool, R. Bates, A. Žídek, A. Potapenko, A. Bridgland, C. Meyer, S. A. A. Kohl, A. J. Ballard, A. Cowie, B. Romera-Paredes, S. Nikolov, R. Jain, J. Adler, T. Back, S. Petersen, D. Reiman, E. Clancy, M. Zielinski, M. Steinegger, M. Pacholska, T. Berghammer, S. Bodenstein, D. Silver, O. Vinyals, A. W. Senior, K. Kavukcuoglu, P. Kohli, D. Hassabis, Highly accurate protein structure prediction with AlphaFold. Nature. 596, 583–589 (2021).

32. M. Zweig, D. Cummings, Cleavage of head and tail proteins during bacteriophage T5 assembly: selective host involvement in the cleavage of a tail protein. Journal of molecular biology. 80 (1973),, doi:10.1016/0022-2836(73)90418-x.

33. L. C. Tsui, R. W. Hendrix, Proteolytic processing of phage lambda tail protein gpH: timing of the cleavage. Virology. 125, 257–264 (1983).

34. S. Kanamaru, P. G. Leiman, V. A. Kostyuchenko, P. R. Chipman, V. V. Mesyanzhinov, F. Arisaka, M. G. Rossmann, Structure of the cell-puncturing device of bacteriophage T4. Nature. 415, 553–557 (2002).

35. S. R. Casjens, R. W. Hendrix, Locations and amounts of major structural proteins in bacteriophage lambda. J Mol Biol. 88, 535–545 (1974).

36. P. Bárdy, T. Füzik, D. Hrebík, R. Pantůček,J. Thomas Beatty, P. Plevka, Structure and mechanism of DNA delivery of a gene transfer agent. Nat Commun. 11, 3034 (2020).

37. L. T. Alexander, R. Lepore, A. Kryshtafovych, A. Adamopoulos, M. Alahuhta, A. M. Arvin, Y. J. Bomble, B. Böttcher, C. Breyton, V. Chiarini, N. B. Chinnam, W. Chiu, K. Fidelis, R. Grinter, G. D. Gupta, M. D. Hartmann, C. S. Hayes, T. Heidebrecht, A. Ilari, A. Joachimiak, Y. Kim, R. Linares, A. L. Lovering, V. V. Lunin, A. N. Lupas, C. Makbul, K. Michalska, J. Moult, P. K. Mukherjee, W. S. Nutt, S. L. Oliver, A. Perrakis, L. Stols, J. A. Tainer, M. Topf, S. E. Tsutakawa, M. Valdivia-Delgado, T. Schwede, Target highlights in CASP14: Analysis of models by structure providers. Proteins. 89, 1647–1672 (2021).

38. G. Guihard, P. Boulanger, L. Letellier, Involvement of phage T5 tail proteins and contact sites between the outer and inner membrane of Escherichia coli in phage T5 DNA injection. J. Biol. Chem. 267, 3173–3178 (1992).

39. I. G. Denisov, S. G. Sligar, Nanodiscs in Membrane Biochemistry and Biophysics. Chem. Rev. 117, 4669–4713 (2017).

40. E. Kandiah, T. Giraud, A. de Maria Antolinos, F. Dobias, G. Effantin, D. Flot, M. Hons, G. Schoehn, J. Susini, O. Svensson, G. A. Leonard, C. Mueller-Dieckmann, CM01: a facility for cryo-electron microscopy at the European Synchrotron. Acta Crystallogr D Struct Biol. 75, 528–535 (2019).

41. X. Li, P. Mooney, S. Zheng, C. R. Booth, M. B. Braunfeld, S. Gubbens, D. A. Agard, Y. Cheng, Electron counting and beam-induced motion correction enable near-atomic-resolution single-particle cryo-EM. Nat. Methods. 10, 584–590 (2013).

42. K. Zhang, Gctf: Real-time CTF determination and correction. Journal of Structural Biology. 193, 1–12 (2016).

43. G. Tang, L. Peng, P. R. Baldwin, D. S. Mann, W. Jiang, I. Rees, S. J. Ludtke, EMAN2: an extensible image processing suite for electron microscopy. J Struct Biol. 157, 38–46 (2007).

44. J. Zivanov, T. Nakane, B. O. Forsberg, D. Kimanius, W. J. Hagen, E. Lindahl, S. H. Scheres, New tools for automated high-resolution cryo-EM structure determination in RELION-3. eLife. 7, e42166 (2018).

45. C.-A. Arnaud, thesis, University Grenoble-Alpes (2017).

46. P. Emsley, B. Lohkamp, W. G. Scott, K. Cowtan, Features and development of Coot. Acta Crystallogr. D Biol. Crystallogr. 66, 486–501 (2010).

47. P. D. Adams, P. V. Afonine, G. Bunkóczi, V. B. Chen, I. W. Davis, N. Echols, J. J. Headd, L.-W. Hung, G. J. Kapral, R. W. Grosse-Kunstleve, A. J. McCoy, N. W. Moriarty, R. Oeffner, R. J. Read, D. C. Richardson, J. S. Richardson, T. C. Terwilliger, P. H. Zwart, PHENIX: a comprehensive Python-based system for macromolecular structure solution. Acta Crystallogr. D Biol. Crystallogr. 66, 213–221 (2010).

48. C. J. Williams, J. J. Headd, N. W. Moriarty, M. G. Prisant, L. L. Videau, L. N. Deis, V. Verma, D. A. Keedy, B. J. Hintze, V. B. Chen, S. Jain, S. M. Lewis, W. B. Arendall, J. Snoeyink, P. D. Adams, S. C. Lovell, J. S. Richardson, D. C. Richardson, MolProbity: More and better reference data for improved all-atom structure validation. Protein Sci. 27, 293–315 (2018).

49. A. P. Pandurangan, M. Topf, RIBFIND: a web server for identifying rigid bodies in protein structures and to aid flexible fitting into cryo EM maps. Bioinformatics. 28, 2391–2393 (2012).

50. R. T. Kidmose, J. Juhl, P. Nissen, T. Boesen, J. L. Karlsen, B. P. Pedersen, Namdinator - automatic molecular dynamics flexible fitting of structural models into cryo-EM and crystallography experimental maps. IUCrJ. 6, 526–531 (2019).

51. M. G. Casabona, Y. Vandenbrouck, I. Attree, Y. Couté, Proteomic characterization of Pseudomonas aeruginosa PAO1 inner membrane. Proteomics. 13, 2419–2423 (2013).

52. J. Cox, M. Mann, MaxQuant enables high peptide identification rates, individualized p.p.b.-range mass accuracies and proteome-wide protein quantification. Nat Biotechnol. 26, 1367–1372 (2008).

53. H. Delattre, O. Souiai, K. Fagoonee, R. Guerois, M.-A. Petit, Phagonaute: A web-based interface for phage synteny browsing and protein function prediction. Virology. 496, 42–50 (2016).

